# Adaptation to visual sparsity enhances responses to infrequent stimuli

**DOI:** 10.1101/2024.03.12.584626

**Authors:** Tong Gou, Catherine A. Matulis, Damon A. Clark

## Abstract

Sensory systems adapt their response properties to the statistics of their inputs. For instance, visual systems adapt to low-order statistics like mean and variance to encode the stimulus efficiently or to facilitate specific downstream computations. However, it remains unclear how other statistical features affect sensory adaptation. Here, we explore how *Drosophila*’s visual motion circuits adapt to stimulus sparsity, a measure of the signal’s intermittency not captured by low-order statistics alone. Early visual neurons in both ON and OFF pathways alter their responses dramatically with stimulus sparsity, responding positively to both light and dark sparse stimuli but linearly to dense stimuli. These changes extend to downstream ON and OFF direction-selective neurons, which are activated by sparse stimuli of both polarities, but respond with opposite signs to light and dark regions of dense stimuli. Thus, sparse stimuli activate both ON and OFF pathways, recruiting a larger fraction of the circuit and potentially enhancing the salience of infrequent stimuli. Overall, our results reveal visual response properties that increase the fraction of the circuit responding to sparse, infrequent stimuli.

## Introduction

Sensory systems adapt to the statistical properties of their inputs by changing their gain, dynamics, and nonlinear properties ^1^. For instance, across sensory modalities, stimuli with higher means tend to reduce the response gain: in olfaction ^2-5^, in audition ^6^, and in vision ^7-10^. Beyond adapting to changing stimulus mean, sensory systems also adapt to the variance, or contrast, in their inputs: this is true in olfaction ^3^, in audition ^11,12^, in somatosensation ^13^, and in visual intensity ^14,15^ and visual motion ^16^. In the cases of both mean and variance, the adaptation tends to shift the neuronal dynamics and dynamic range so that they can efficiently encode the stimulus ^16-19^, but the changes may also be seen in light of specific downstream computations ^20,21^. Though sensory systems also adapt to some types of higher order stimulus statistics ^22-25^, it remains unclear how the adaptation-driven changes affect downstream computations. In this study, we examine the intuitive statistic of visual stimulus sparsity and investigate how processing changes driven by stimulus sparsity affect the downstream canonical computation of visual motion.

In the visual system, early neurons adapt strongly to both stimulus mean and variance. In insects and vertebrate eyes, photoreceptors adapt strongly to the mean of the stimulus intensity and then report fractional deviations away from that mean ^8,10,26-29^. Downstream from photoreceptors, in both insects and vertebrates, neurons adapt to brighter stimuli by narrowing their receptive fields in space and time and passing less signal at low frequencies ^17,19,30,31^, as well as by changing other nonlinear properties ^32,33^. Both insects and vertebrate visual systems adapt to the variability in intensity, or contrast, of stimuli ^14,15,20,21^. This adaptation tends change gains so that visual neurons respond strongly to the full range of the presented stimulus, but can also change the response dynamics ^14,21,34-36^. Vertebrate retina also adapts to a range of other spatiotemporal stimulus correlations ^37^.

Beyond the stimulus mean and variance, adaptation to higher-order stimulus statistics has also been investigated in vertebrate visual systems. There, the stimulus skewness and kurtosis, the third and fourth moments of the stimulus distribution, do not induce substantial changes in response properties ^22,23^. But the phase correlations of stimuli, related to their spatial structure, induce substantial adaptation in cortical visual neurons ^24,25^. However, phase correlations do not capture all spatial stimulus features, and sensitivity to phase correlations does not always have intuitive interpretations or relationships with the processing goals of the visual system. In comparison, here we examine the stimulus statistic of sparsity — how frequently stimulus values are nonzero over space or time. Sparsity is an intuitive feature that is not easily captured by moments of the stimulus distribution. The sparse nature of many natural signals is critical to signal compression and reconstruction ^38,39^, and is thus a natural feature for sensory system sensitivity. In theories of sensory coding, sparsity is related to the concept of “sparse coding”, a strategy that encodes sensory information in the activity of only a small fraction of neurons ^40-42^.

Here, we have investigated how circuits adapt to changes in stimulus sparsity. To do this, we focused on motion detection circuitry in the fruit fly *Drosophila* because its anatomy and computational structure are well studied ^43^, while powerful genetic tools allow us to measure and manipulate its activity. We begin by showing that natural scenes exhibit many different degrees of sparsity. To characterize how visual neurons change their responses to stimuli with different sparsity, we measured calcium responses to parameterized sparse and dense visual stimuli, showing how timecourses change dramatically with stimulus sparsity (dense vs. sparse) and polarity (light vs. dark), but are distinct from the phenomenon of contrast adaptation. A phenomenological linear-nonlinear-linear (LNL) model reproduced much of the sparsity adaptation phenomenology, but optogenetic experiments ruled out that the effects are due to simple calcium accumulation. Calcium measurements revealed widespread sparsity adaptation across visual neurons in both the ON and OFF pathways. In both ON and OFF pathways, dense stimuli lead to responses that change sign with stimulus polarity, while sparse stimuli yield activation independent of stimulus polarity. By measuring receptive fields, we showed that sparsity adaptation exerts a pronounced impact on the receptive field center but not on its surround. To investigate how the observed processing changes affect motion detection, we probed visual system responses using sparse and dense moving bars. These stimuli yielded strong adaptation to sparsity, including in the primary direction-selective cells, which each responded to both light and dark isolated moving bars but with different tunings. These experiments show that sparse ON and OFF stimuli each elicit positive responses in both ON and OFF pathway circuits, including in motion detectors, while dense moving bars elicit opposite responses in ON and OFF pathways. Thus, stimuli with different sparsity alter visual processing to recruit stronger and broader neural responses to infrequent stimuli.

## Results

### Sparsity varies within natural scenes

Intuitively, a sparse signal is one in which the non-zero coefficients only occupy a very small portion of its representation ^45^. Since real signals vary in their amplitude, a measure of sparsity should not depend on the absolute amplitude of the signal but should rather reflect the magnitude differences among coefficients. We developed a sparsity index that describes the relative distribution of contrast in a patch of visual space, assuming that a small dark object is at its center (see **Methods** for details). A sparse signal is one in which most contrasts are near 0 except for the dark spot; such a signal would yield a sparsity index near 0. Conversely, a dense signal has many contrasts far from 0 and a sparsity index near 1. We used this metric to characterize natural scenes (**Fig. 1A**). Sparse regions predominantly occur in the sky where there is mostly uniform luminance. There, a small dark object against the sky contains almost all the contrast power. On the other hand, dense regions exist in cluttered areas of the foreground, where there are large local luminance fluctuations. When we plotted the distribution of local sparsity at three different spatial scales over many images (**Fig. 1B**), it revealed significant variability in local sparsity across natural scenes. Importantly, this spatial variability in sparsity extends to temporal variability when a sighted organism moves within a natural scene and the scene moves across its retina. The natural spatial and temporal variability in sparsity motivated our investigation into visual responses to stimuli with varying degrees of sparsity.

**Figure 1.**
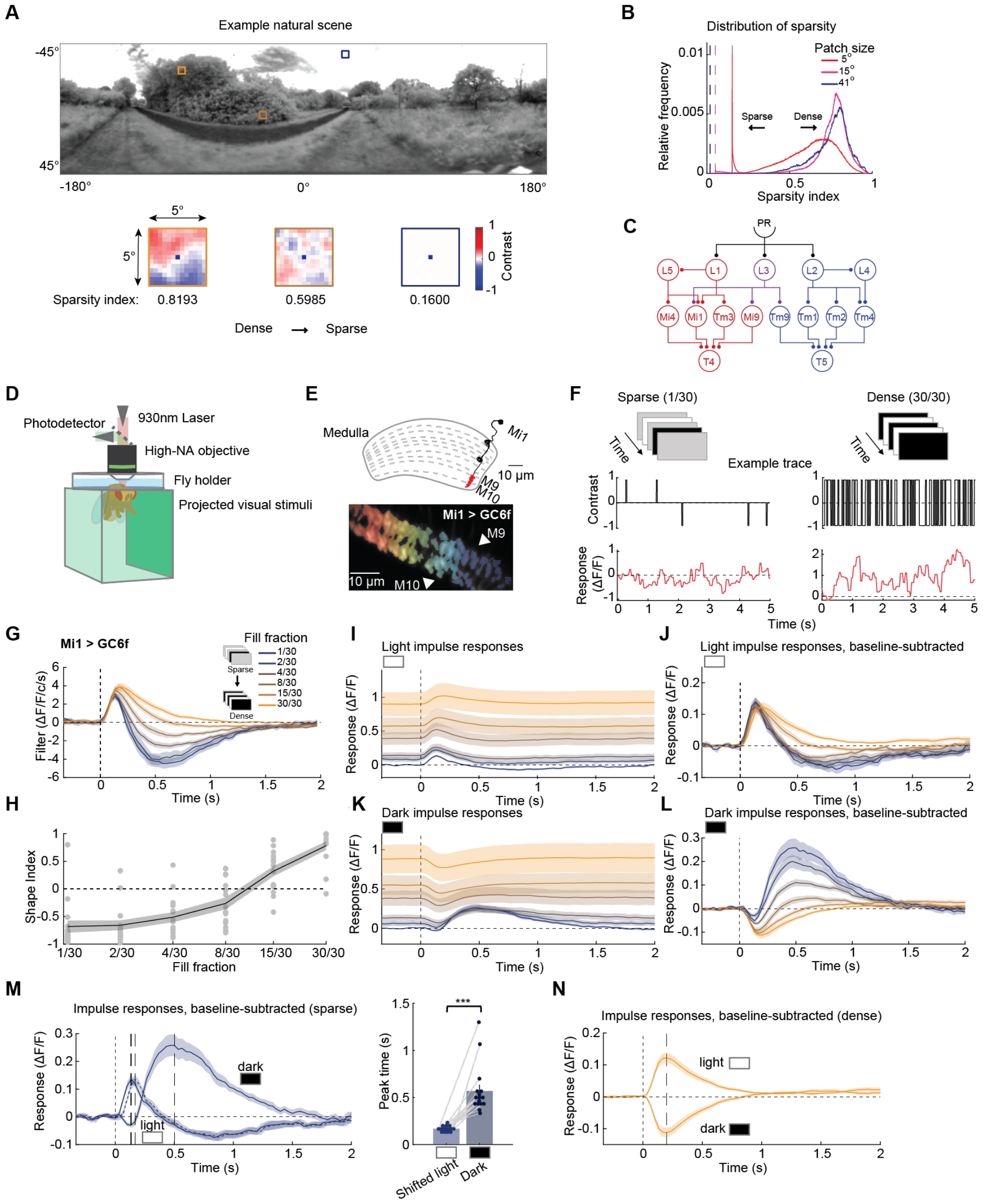
Variation of sparsity in natural scenes and strong sparsity adaptation in Mi1 neurons. (A) A natural scene from the database ^44^ with three sample regions exhibiting different sparsities. Local sparsity was computed as described in the methods. (B) Distribution of the calculated sparsity indices over many natural scenes. (C) Circuit diagram showing a subset of feedforward elementary motion detection neural circuitry in *Drosophila*. (D) Schematic of the two-photon microscopy and panoramic screen setup. (E) Mi1 neuronal anatomy. *Top*: Mi1 has dendrites in medulla layers M1 and M5, while its axon terminals are in medulla layers M9 and M10. *Bottom*: image of axon terminals of Mi1 neurons expressing GCaMP6f. Color coded regions of interest (ROIs) in the axon terminal layers. (F) *Top*: Schematic of sparse (left) and dense (right) ternary noise visual stimulus; stimulus frames are updated at 30 Hz. *Bottom*: Example time traces of visual stimulus and recorded calcium responses. (G) Linear filters extracted to different fill fraction (sparsity) epochs (N = 16 flies). (H) Shape index calculated for filters computed from different sparsity stimuli. (I) Light impulse responses to stimulus epochs with different fill fraction (sparsity). (J) Baseline subtracted light impulse responses to stimulus epochs with different fill fraction (sparsity). (K) Dark impulse responses to stimulus epochs with different fill fraction (sparsity). (L) Baseline subtracted dark impulse responses to stimulus epochs with different fill fraction (sparsity). (M) Left: baseline-subtracted light and dark impulse responses overlaid for the sparsest stimulus. A light impulse response shifted by one frame was also added to compare the peak times of light and dark impulse responses. Right: Comparison of shifted light impulse response peak time with dark impulse response peak time. (N) Baseline-subtracted light and dark impulse responses overlaid for 30/30 fill fraction (least sparse case). Error bars and shading around mean traces all indicate standard error of the mean. n. s.: not significant (p > .05); *: p < .05; **: p < .01; ***: p < .001; ****: p < .0001 in Wilcoxon signed-rank tests across flies.

### Mi1 calcium measurements show strong sparsity adaptation

To investigate how neurons respond to sparse and dense visual stimuli, we focused on the fly’s motion detection circuitry. There, photoreceptors detect light intensity and transmit the information to lamina cells L1-L3 (**Fig. 1C**). Signals are then split into two parallel pathways: an ON pathway encoding luminance increments and an OFF pathway encoding luminance decrements ^46-48^. Then, at least four different types of these ON and OFF medulla cells synapse onto the direction-selective (DS) neurons T4 and T5 ^49^, which are required for optomotor rotational responses ^50^ and optomotor speed regulation ^51^. We first expressed GCaMP6f ^52^ in Mi1 neurons, ON neurons that are required for T4 function ^47,53^. We then measured fluorescence responses in Mi1 elicited by a ternary noise stimulus (**Fig. 1D-F**), in which the temporal sparsity could be modulated by changing the fraction of total frames that were light/dark flashes (fill fraction) (see **Methods**).

From the responses of Mi1 neurons to this stimulus (**Fig. 1F**), we first extracted linear filters to describe the dynamic responses to stimuli with different sparsities (see **Methods**). The shape and amplitude of the filters changed substantially as the fill fraction increased and the stimulus became denser (**Fig. 1G**). We quantified these changes with the shape index (**Fig. 1H**, see **Methods**), which measures the fraction of the filter area that is in the positive lobe versus negative lobe. When the input signal was sparse, the filter has a prominent negative lobe, with shape index close to -1. However, as the fill fraction increases, the filter becomes less biphasic and the shape index approaches 1, and the amplitude also slightly increases. These findings show that the temporal filtering properties of Mi1 depend on sparsity and cannot be fully described by a single linear filter. In principle, changing the input statistics alone could result in different filter estimates without adaptation occurring ^54^; however, a simple linear-nonlinear model does not show changing filters as the stimulus sweeps from sparse to dense stimuli (**Fig. S1A-B**). In this study, we will define adaptation as a change in linear or nonlinear processing properties of a cell, a standard definition used in many past studies ^1,14,15,23^. The change in the linear filter shape is therefore a hallmark of adaptation.

We next asked how sparsity affects Mi1 responses to light and dark stimuli, since our stimulus contains both light and dark flashes. We calculated the light and dark impulse responses (**Fig. 1I, K**) for each fill fraction by computing the mean response triggered by either light or dark flashes (see **Methods** for more details). The baseline calcium signal increased with increasing fill fraction, indicating accumulated calcium during stimulus presentation, consistent with previous findings ^21^. To examine the effect of sparsity beyond baseline shifts, we subtracted the baselines from all the impulse responses (**Fig. 1J, L**). In the sparse condition, light impulses initially increased the calcium activity and subsequently suppressed it in Mi1 cells, while dark impulses elicited a small initial suppression followed by a large activation, consistent with prior measurements ^55^. Importantly, the delay in the peak of the dark impulse response lagged far behind the light impulse response peak, even after aligning the light impulse response with the trailing luminance increment (**Fig. 1M**). Therefore, the response to the dark flash is not a simple response to the luminance increment as the stimulus returned to baseline. Conversely, in the dense condition, the impulse responses were symmetrical about the baseline activity, with light impulses resulting in more activity and dark impulses resulting in less (**Fig. 1N**). This symmetry is imposed by the structure of the dense, binary stimulus (see **Methods, Fig. S1C-E**). However, even when the symmetry imposed by the dense stimulus was removed by using a uniform distribution of luminance levels, responses to sparse and dense stimuli were still markedly different and the responses to the dense stimulus still look predominantly linear (**Fig. S1F-H**). This demonstrates that the response differences measured to sparse and dense stimuli are not due to the structure of our dense stimulus and represent sparsity adaptation within Mi1.

**Supplementary Figure S1.**
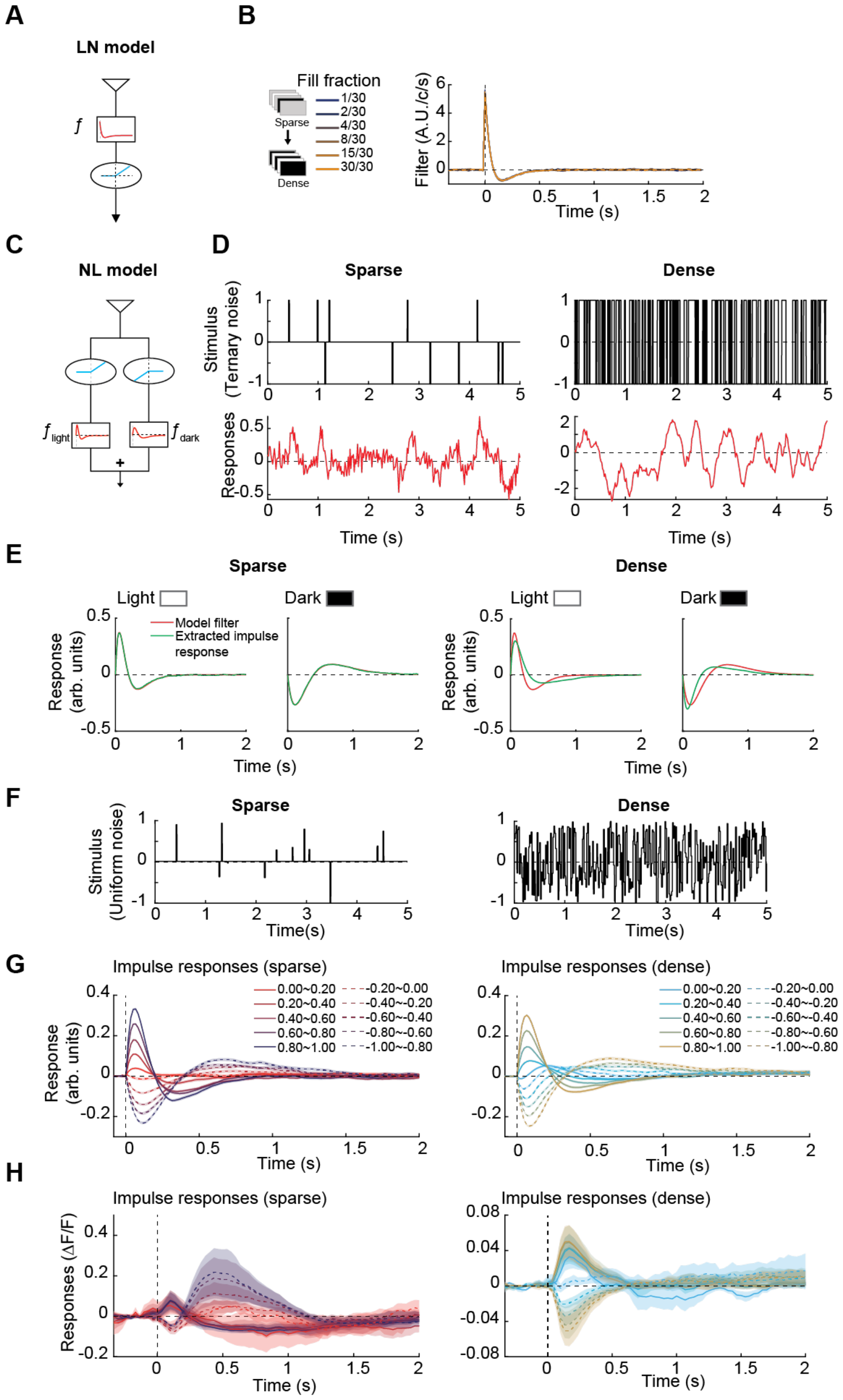
Impact of stimulus on impulse response measurements. (Related to Figure 1) (A) Simple linear-nonlinear (LN) model. (B) Extracted linear filters from the LN model in response to varying stimulus sparsity. Stimulus sparsity does not alter the filter shape for this model; not all line colors are visible because the extracted filters fall exactly on top of one another. (C) Nonlinear-linear (NL) model that creates distinct responses to light and dark impulses. (D) Example time traces of the ternary noise stimuli and NL model output responses. (E) Extracted light and dark impulse responses to sparse (left) and dense (right) stimulus overlaid with the original filters of the NL model. In the sparse case, the calculated impulse responses aligned with the original NL model filters. However, the dense, binary stimulus generates equal and opposite light and dark impulse responses (see **Methods**), diverging from the original filters. (F) Example time traces of sparse (left) and dense (right) stimuli with contrasts drawn from a uniform distribution. (G) Extracted impulse responses corresponding to various contrast bins from the responses of the NL model to the sparse (left) and dense (right) uniform noise stimuli. Here, the dense impulse responses in the NL model no longer appear equal and opposite and effectively regained the original NL model filters. (H) Extracted impulse responses corresponding to various contrast bins from the Mi calcium responses to the sparse (left) and dense (right) uniform noise stimuli. (N = 7 flies). Here, the dense stimuli still elicit light and dark impulse responses that are close to equal and opposite, and importantly, very different in shape from those elicited by sparse stimuli. Error bars and shading around mean traces all indicate standard error of the mean.

### Sparsity adaptation is distinct from contrast adaptation

When a stimulus becomes less sparse, the contrast of the stimulus, measured as the root-mean-squared deviation from the mean, also becomes larger. Thus, we asked whether sparsity adaptation was simply the same phenomenon as contrast adaptation, which has been characterized in both vertebrate retinae ^14,15,56^ and in the fly visual system ^20,21^. Both in contrast adaptation and in responses to stimuli with different sparsity (**Fig. 1**), there are substantial changes in baseline as stimuli become more dense/higher contrast ^14,21^. To examine how sparsity adaptation relates to contrast adaptation, we systematically varied the stimulus contrast in both sparse and dense stimuli and measured Mi1 responses. Irrespective of stimulus contrast, the extracted linear filters for sparse and dense stimuli had distinct shapes, with sparse stimuli eliciting filters with a prominent negative lobe and dense stimuli eliciting monophasic filters (**Fig. 2A, B**). Thus, the two separate temporal filtering regimes are independent of contrast. As before, we plotted the shape index of the filters for sparse and dense stimuli, which revealed two distinct shapes, consistent over contrast values (**Fig. 2C**). In the dense regime, the amplitude of filters varied over contrast, consistent with earlier measures of contrast adaptation (**Fig. S2B**) ^21^. Higher contrast yielded higher baseline responses as well (**Fig. S2C-E**). Like the filters, light and dark impulses responses to sparse and dense stimuli both have consistent shapes across the tested contrast range (**Fig. 2D, E, S2D, S2E**). Therefore, the shape changes associated with different sparsity stimuli are specific to sparsity itself and are not a result of changes in contrast. Last, to check whether the observations depended on the calcium indicator, we measured the calcium response of Mi1 to the same ternary noise stimulus using a different calcium indicator, GCaMP7b ^57^ (**Fig. S2F-L)**. These results aligned closely with the earlier results (**Fig. 1**), indicating they are not strongly dependent on the indicator.

**Figure 2.**
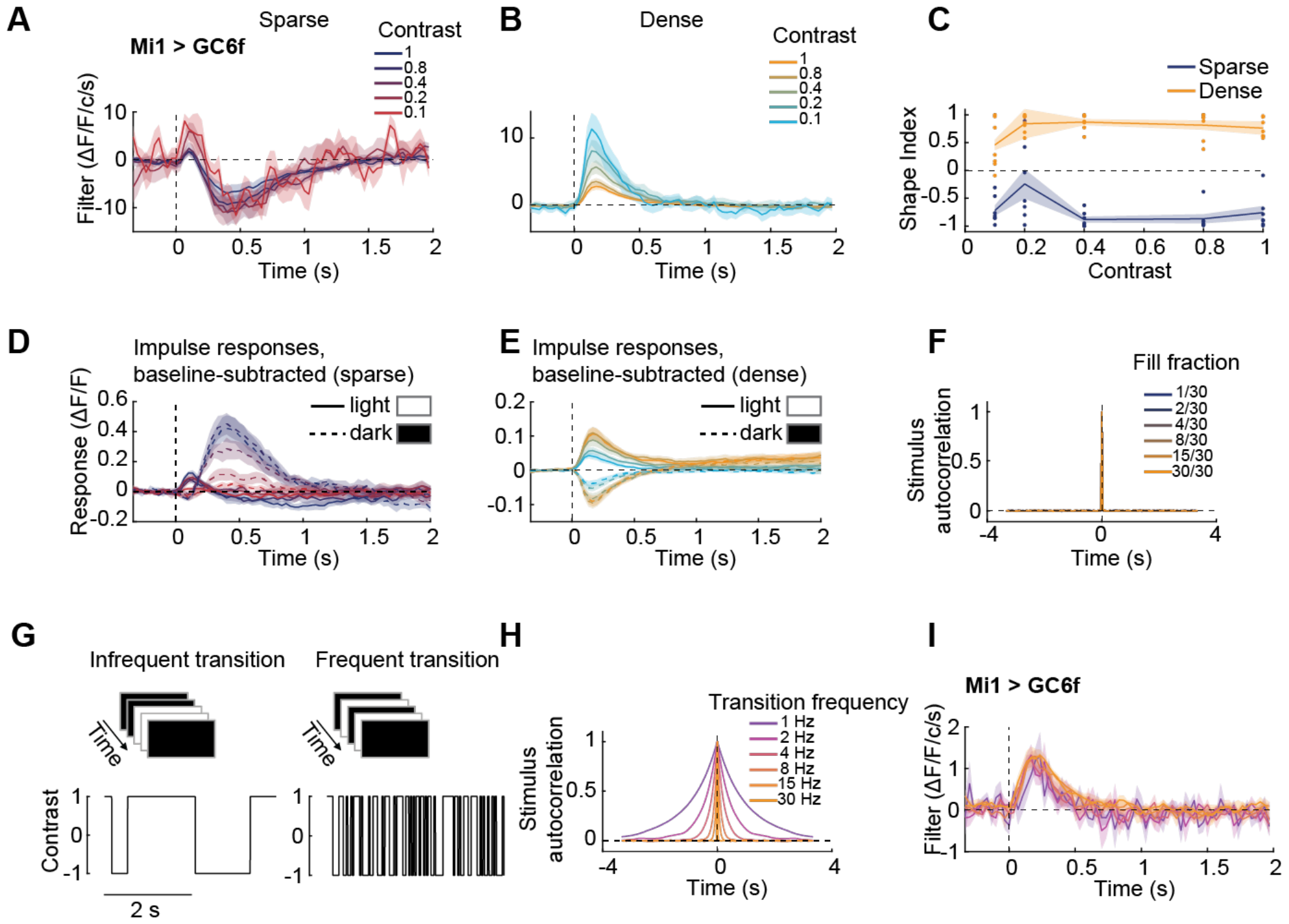
Characterizing sparsity adaptation. (A, B) Linear filters extracted to sparse (A) and dense (B) stimuli in which stimuli’s amplitude were swept from 0.1 to 1 to change the contrast (left). (N = 7 flies). (C) Shape index calculated for each filter of different contrast amplitudes. (D, E) Baseline-subtracted light (solid lines) and dark (dashed lines) impulse responses to sparse (D) and dense (E) stimuli. (F) Signals’ autocorrelations for each epoch in the ternary noise stimulus. (G) Correlated version of the ternary noise stimulus. (H) Signals’ autocorrelations for each epoch in the correlated stimulus. (I) Linear filters extracted to each epoch from the responses to the correlated stimulus. (N = 8 flies). Error bars and shading around mean traces all indicate standard error of the mean.

**Supplementary Figure 2.**
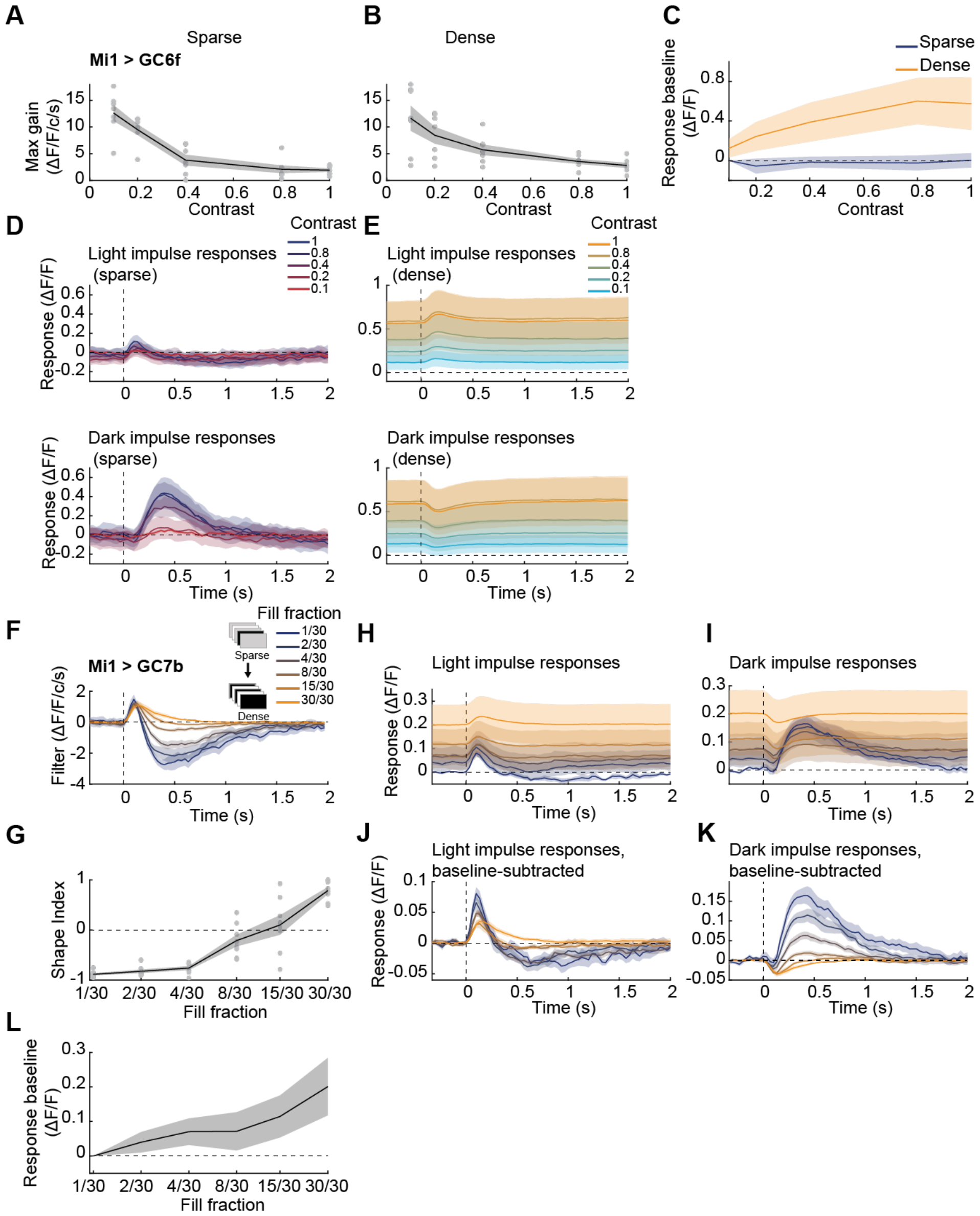
Contrast changes the basal calcium response and filter gain in Mi1 and calcium responses in Mi1 measured with GC7b. (Related to Figure 2) (A, B) Filters’ gain corresponding to each contrast amplitude in sparse (A) and dense (B) conditions. (C) Basal calcium response level to the sparse and dense stimuli along with the contrast sweep. (D, E) Extracted light and dark impulse responses to sparse (left) and dense (right) stimulus. (F-L) Calcium measurements in Mi1 using GC7b: linear filters (F), shape index (G), light (H) and dark (I) impulse responses, response baseline change (L) and baseline subtracted light (J) and dark (K) impulse responses corresponding to stimuli with different fill fractions (sparsity). (N = 8 flies). Error bars and shading around mean traces all indicate standard error of the mean.

### Sparsity adaptation depends on contrast excursions, not contrast transitions

Retinal ganglion cells adjust filtering properties based on temporal correlations in the input signals ^37^. The dense and sparse stimuli we use are uncorrelated in time (**Fig. 2F**), so the adaptation cannot be related to temporal correlation structure per se. However, in our sparse stimuli, changes in the stimulus occur infrequently, by construction. Could it be that cells are adapting to frequency of changes in the stimulus, rather than to the sparsity of deviations from 0 contrast? To answer this question, we designed a dense stimulus in which all frames were not gray, but in which contrast changed with different frequencies. This amounts to a binary stimulus with intensities that correlate over time (**Fig. 2G**). In this stimulus, the transition frequency was systematically varied from 1 Hz to 30 Hz, so that the transitions in this stimulus matched the rate of the deviations from gray in our original stimulus. However, this new stimulus is considered dense by our metrics, since its fill fraction is always 30/30. The autocorrelation in contrast of these stimuli is an exponential decay in time (**Fig. 2H**). The linear filters extracted from responses to these stimuli had similar shapes over the range of transition frequencies tested (**Fig. 2I**), indicating that Mi1 neurons adapt to the fraction of light/dark states themselves rather than to the frequency of transitions between states.

**Supplementary Figure 3.**
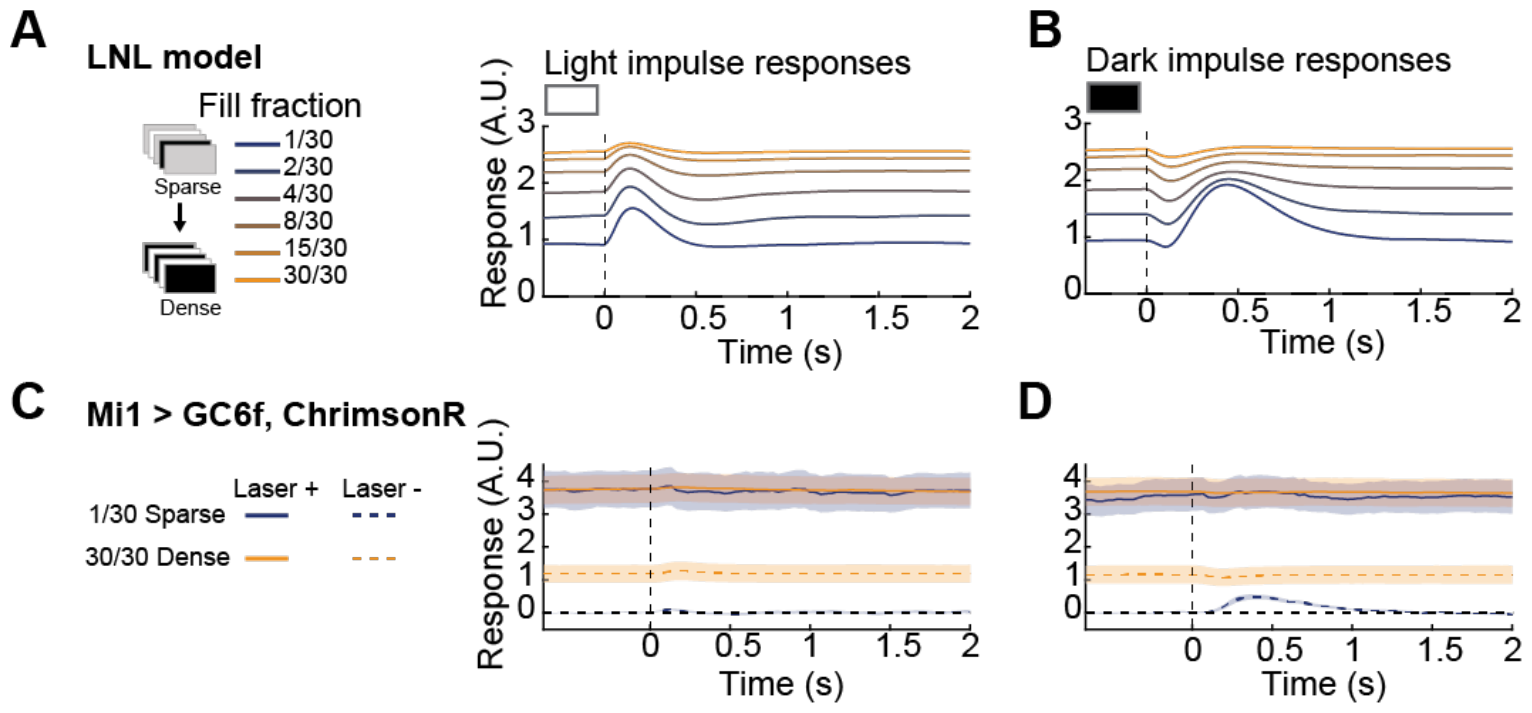
Additional results from the computational model simulation and optogenetic activation of Mi1. (Related to Figure 3) (A, B) Light (A) and dark (B) impulse responses calculated from the LNL model response to the ternary noise stimulus. (C, D) Light (C) and dark (D) impulse responses before and after optogenetic activation of Mi1 for both sparse and dense stimuli. (N = 7 flies).

### A biophysically plausible model replicates phenomenology of sparsity adaptation

To investigate potential mechanisms responsible for sparsity adaptation, we developed a biophysical plausible linear-nonlinear-linear (LNL) model with fixed parameters that was able to replicate the observed phenomenon. In this model, stimuli are filtered by a high pass filter before a saturating point nonlinearity is applied and the resulting signal is filtered by a second filter, this time a low pass filter (**Fig. 3A**). This model is similar to ones previously analyzed as a source for contrast adaptation ^21,58^, but here we changed the nonlinearity to rectify the inputs. In this type of model, the two linear filters can effectively ‘hide’ the rectifying and saturating nonlinearity, making it difficult to observe directly ^21^. This also belongs to a class of models that can exhibit complex adaptation phenomena without model parameters changing over time ^58-60^.

**Figure 3.**
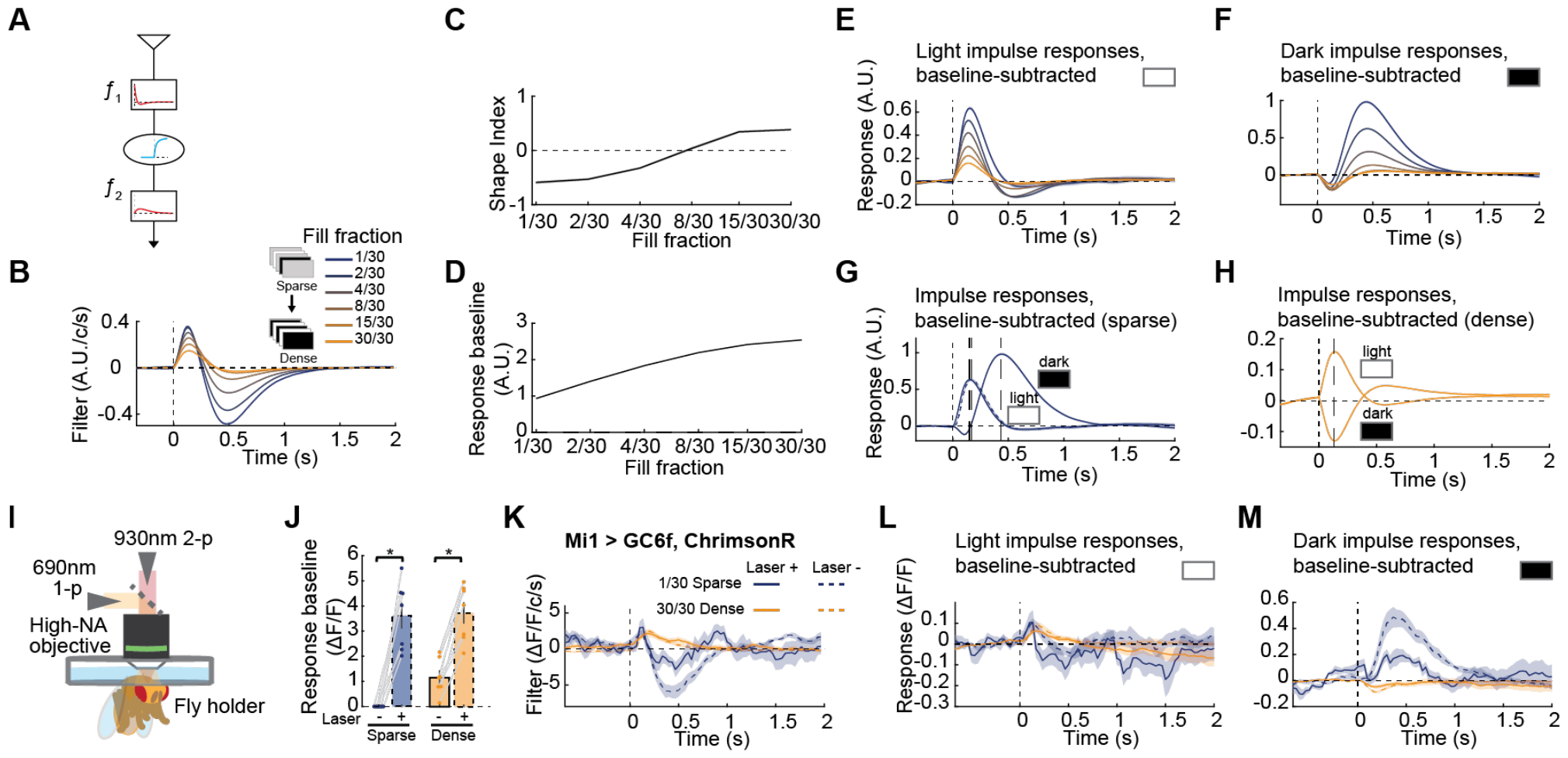
A phenomenological model for sparsity adaptation in Mi1. (A) A biophysical plausible linear-nonlinear-linear (LNL) model can reproduce many of the sparsity adaptation phenomena in Mi1. (B-H) Analysis results from the LNL outputs in response to varying stimulus sparsity: linear filters (B), shape index (C), baseline increases (D), baseline subtracted light (E) and dark (F) impulse responses, baseline-subtracted light and dark impulse responses overlaid for the sparsest (G) case and densest cases (H). (I) Schematics of the setup for the optogenetic activation during two-photon imaging experiments. (J) Calcium response baselines before and after optogenetic activation of Mi1 for both sparse and dense stimuli. (K-M) Comparison of Mi1 responses to sparse and dense stimuli with and without optogenetic activation: linear filters (K), baseline subtracted light (L) and dark (M) impulse responses. (N = 7 flies). Error bars and shading around mean traces all indicate standard error of the mean. n. s.: not significant (p > .05); *: p < .05; **: p < .01; ***: p < .001; ****: p < .0001 in Wilcoxon signed-rank tests across flies. Error bars and shading around mean traces all indicate standard error of the mean.

This model successfully replicates the filter shape changes as stimuli transition from sparse to dense (**Fig. 3B, C**). The model also successfully replicates the calcium accumulation, where the baseline response increased with denser stimuli (**Fig. 3D, S3A, B**). Moreover, the shape of the model impulse responses varied with stimulus sparsity (**Fig. 3E, F, S3A, B**), with patterns similar to those measured in Mi1 (**Fig. 1**). Under sparse conditions, light impulses elicit in the model an initial activation followed by a subsequent suppression. Conversely, dark impulses elicit a small suppression followed by a large subsequent activation. Furthermore, the pronounced delay in the peak of the dark impulse response persists, relative to the peak of the shifted light impulse response aligned with the trailing luminance increment (**Fig. 3G**). In contrast, under dense stimulus conditions, the model’s light and dark impulse responses were approximately symmetric around the baseline (**Fig. 3H**). Hence, this LNL model replicated both filter and impulse response properties of sparsity adaptation in Mi1. This model is biophysical plausible since many signal processing steps result in filtering in time, and many biological nonlinearities saturate. For example, the saturation could occur at presynaptic neurotransmitter release, postsynaptic neurotransmitter receptors, receptor-gated ion channels, membrane voltage, or voltage-gated calcium channels. This suggests that it may be straightforward to generate the phenomenology measured here in many ways in different neurons and circuits.

### Optogenetic stimulation of Mi1 reveals adaptation occurs upstream of Mi1 voltage

We hypothesized that the saturation step in the LNL model might correspond to a saturation in the calcium flux with respect to voltage, while the low pass filter might correspond to the low passing of the signal due to calcium efflux. To test this hypothesis, we optogenetically activated Mi1 neurons to examine whether constitutively depolarizing Mi1 can affect the shape of response filters or impulse responses. To do this, we simultaneously expressed the fluorescent calcium indicator GCaMP6f and a red-shifted channelrhodopsin Chrimson ^61^ in Mi1 neurons and recorded the fluorescence responses of Mi1 when flies were presented the ternary noise stimuli. A 690-nm laser was shone onto the fly’s brain to activate the Chrimson expressed in Mi1 neurons (**Fig. 3I**). The laser intensity was calibrated so that Mi1 neuron calcium baselines increased by ∼300% in these experiments (**Fig. 3J**). We compared the stimulus filters when Mi1 neurons were optogenetically activated to when they were not, and found that the shapes remained largely unchanged (**Fig. 3K**). Specifically, the filters were biphasic under sparse stimuli, but became monophasic as the stimulus became dense. Likewise, constitutively depolarizing Mi1 neurons did not transform the shape of the sparse impulse responses into the shape of dense impulse responses (**Fig. 3L, M**). These results demonstrate that sparsity adaptation is not strongly impacted by the constitutive optogenetic activation. Thus, sparsity adaptation is not a result of calcium accumulation in Mi1 or constitutive depolarization of the neuron; instead, the adaptation likely occurs in signals upstream of the optogenetic activation.

### Sparsity adaptation occurs broadly across the motion detection circuit

Since sparsity adaptation appears to occur upstream of Mi1 calcium activity (**Fig. 3**), we set out to characterize how broad this adaptation is across *Drosophila*’s motion detection circuitry. We began by examining signals upstream of the calcium signal in Mi1. We expressed the Arclight voltage indicator ^62^ in Mi1 to measure the membrane potential responses of Mi1 to stochastic stimuli with variable sparsity (**Fig. 4B**). Since voltage signals are faster than calcium signals, we extracted filters using a method to obtain higher temporal resolution than the imaging framerate ^63^. Interestingly, the voltage responses were similar to the calcium responses: the voltage filters transitioned from biphasic to monophasic as the stimulus changed from sparse to dense (**Fig. 4Bii**). In both the sparse and dense cases, light and dark voltage impulse responses had much the same shape as the calcium impulse responses, though they were faster in time, as expected (**Fig. 4Biii, iv**). This demonstrates that sparsity adaptation is already present in Mi1 voltage signals.

**Figure 4.**
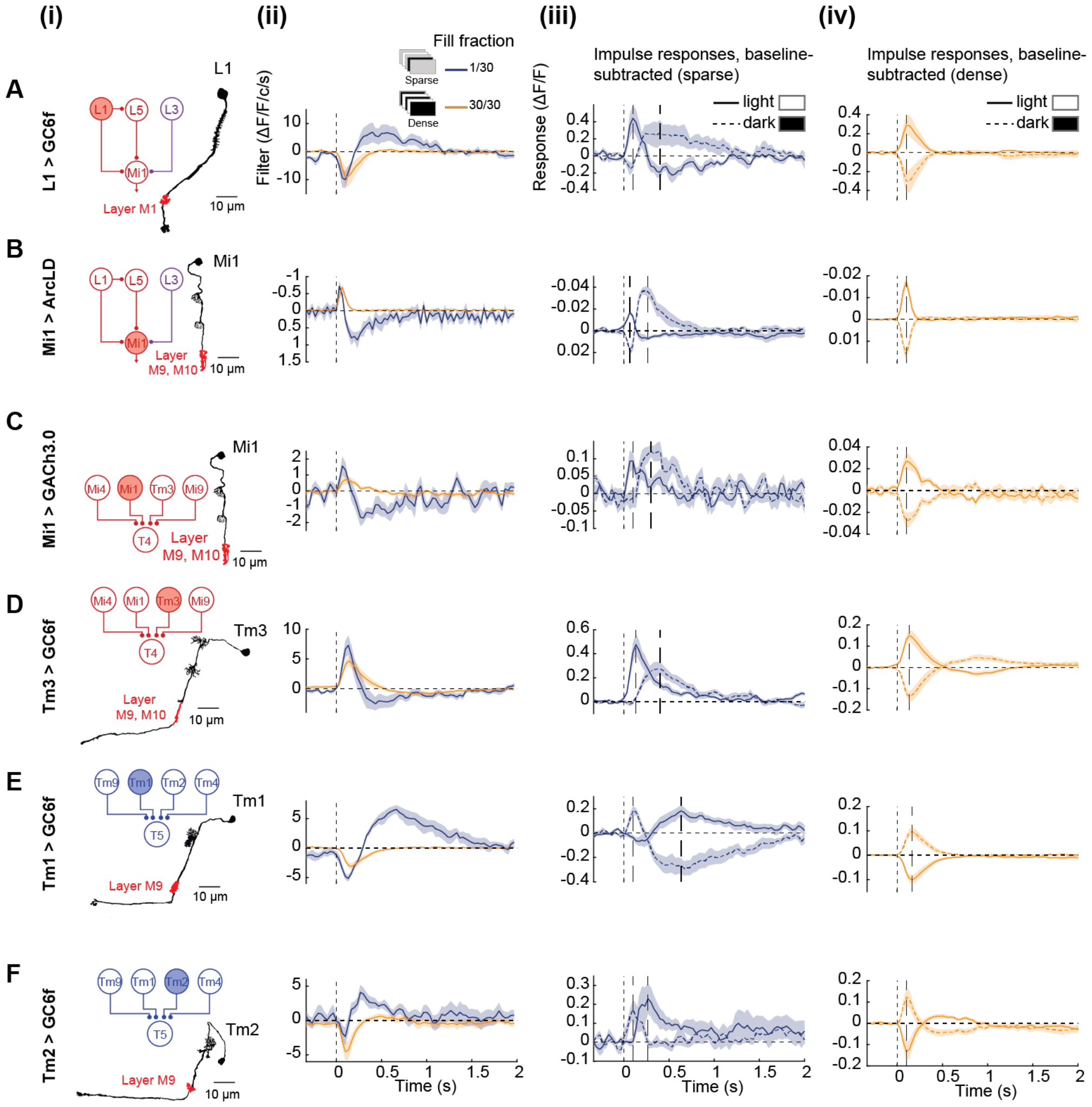
Sparsity adaptation exists widely in motion detection circuits. (A) (i) Summary circuit diagram and neural anatomy depicting neurons within motion detection circuits and where signals were measured for L1 intracellular calcium. (ii) Linear filters extracted to sparse and dense stimuli. (N = 7 flies). (iii, iv) Baseline-subtracted light and dark impulse responses to the sparse (iii) and dense (iv) stimuli. (B) As in (A), except signals were measured for Mi1 membrane voltage. (N = 7 flies). (C) As in (A), except signals were measured for extracellular acetylcholine (ACh) at Mi1 axons. (N = 8 flies). (D) As in (A), except signals were measured for Tm3 intracellular calcium. (N = 11 flies). (E) As in (A), except signals were measured for Tm1 intracellular calcium. (N = 9 flies). (F) As in (A), except signals were measured for Tm2 intracellular calcium. (N = 7 flies). Error bars and shading around mean traces indicate standard error of the mean.

Next, we examined calcium and neurotransmitter signals upstream of Mi1. Mi1 neurons have 3 main inputs: L1, L3, and L5, of which L1 and L3 are directly postsynaptic to photoreceptors, while L5 receives input mainly from L1 ^64,65^. L1 and L3 neurons respond to contrast decrements, with L3 responding more slowly and signaling long term changes in luminance ^55,66,67^. L5 neurons respond transiently to contrast increments ^21^. To test whether these neurons adapt to stimulus sparsity, we measured calcium responses in these neurons to our stimuli with varying sparsity. L1 and L5 showed strong sparsity adaptation, their filters changing from biphasic to monophasic as the stimulus changed from sparse to dense (**Fig. 4A, S4B**). Impulse responses in these neurons show a strong qualitative similarity to those in Mi1. On the other hand, L3 filters show minimal shape change when the stimulus changes from sparse to dense (**Fig. S4A**), indicating that L3 does not adapt to stimulus sparsity. This may align with its proposed functional role as a luminance-sensitive neuron ^66^. To examine another signal upstream of Mi1, we measured extracellular glutamate concentration at Mi1 dendrites (**Fig. S4C**), presumably released primarily by L1 ^21,68^. The glutamate signals at Mi1 dendrites showed some sparsity adaptation, but with less pronounced shape changes in the filters and impulse responses as the stimulus transitioned from sparse to dense (**Fig. S4Cii-iv**). This could differ from the L1 calcium signal if the glutamate signal also includes signals from Dm or other widefield cells ^69^.

To discover whether the sparsity adaptation measured in Mi1 calcium signals translated into Mi1 synaptic release, we expressed GACh3.0 ^70^ in Mi1 to measure extracellular acetylcholine concentrations at its synapse onto T4 (**Fig. 4C**) ^65^. The filters and impulse responses in the acetylcholine signals largely mirrored those in Mi1 calcium signals. This indicates that Mi1’s adaptation is passed on to downstream neurons through neurotransmitter release.

To better understand how sparsity adaptation might affect direction selective circuitry, we next sought to characterize sparsity adaptation in several neurons parallel to Mi1 in the motion detection circuits, which constitute a subset of required input neurons to motion detectors T4 and T5 ^53,71^. We measured sparsity adaptation in Tm1, Tm2, and Tm3, covering ON and OFF excitatory inputs to T4 and T5. Tm3 neurons adapt to stimulus sparsity, although the filter shape change is not as pronounced as Mi1 (**Fig. 4D**). In the OFF pathway, both Tm1 and Tm2 neurons adapt to stimulus sparsity, although Tm1 shows stronger filter shape changes than Tm2 (**Fig. 4E, F**). All three neurons showed delayed activation to non-preferred contrast impulses, consistent with prior measurements ^55^. These results demonstrate widespread sparsity adaptation in motion detection circuits.

**Supplementary Figure 4.**
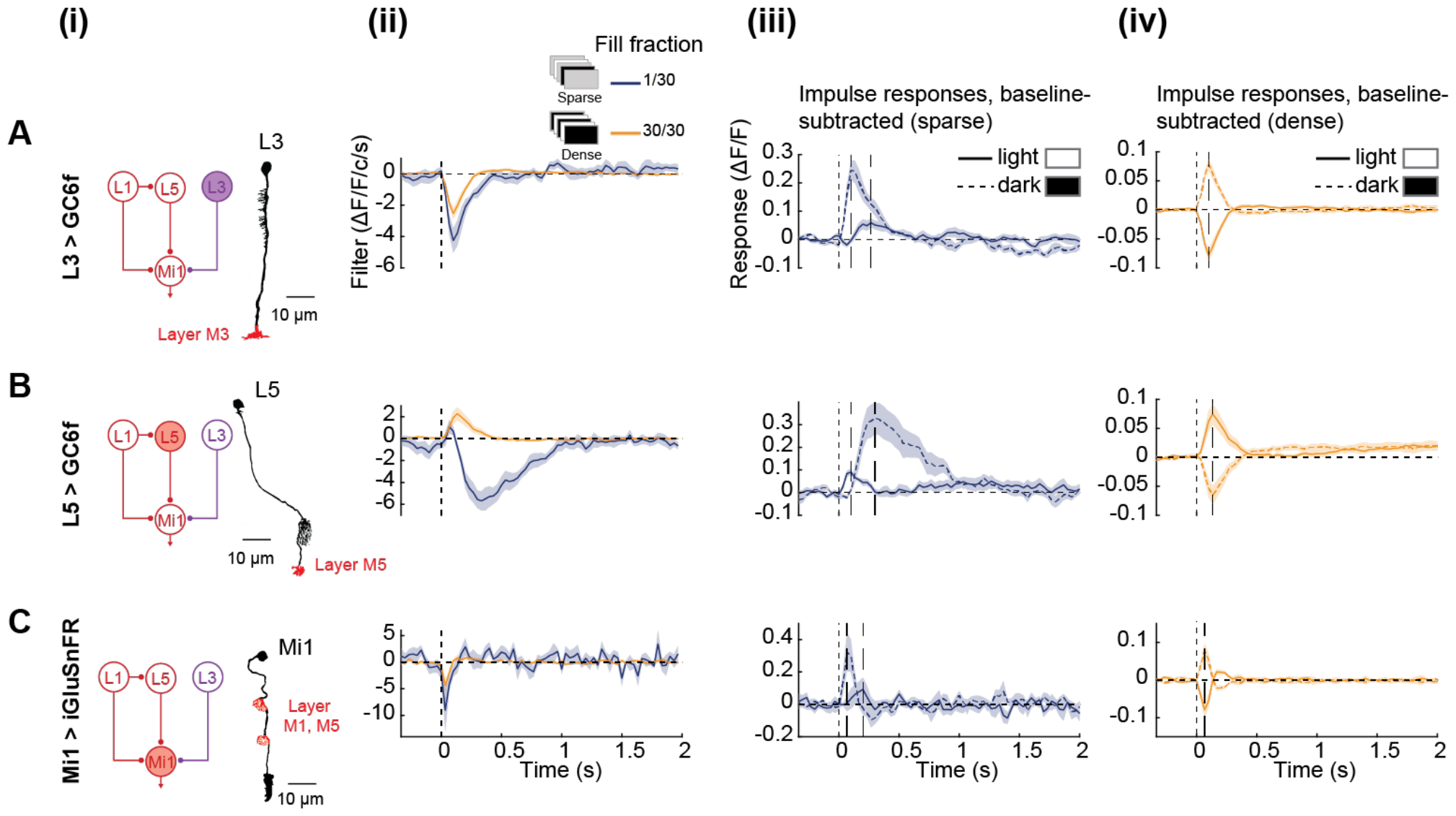
Additional measurements on neurons upstream of Mi1 in the motion detection circuits. (Related to Figure 4) (A) (i) Summary circuit diagram and neural anatomy depicting neurons within motion detection circuits and where signals were measured for L3 intracellular calcium. (ii) Linear filters extracted to sparse and dense stimuli. (N = 8 flies). (iii, iv) Baseline-subtracted light and dark impulse responses to the sparse (iii) and dense (iv) stimuli. (B) As in (A), except signals were measured for L5 intracellular calcium. (N = 9 flies). (C) As in (A), except signals were measured for extracellular glutamate (Glu) at Mi1 dendrites. (N = 7 flies). Error bars and shading around mean traces indicate standard error of the mean.

**Supplementary Figure 5.**
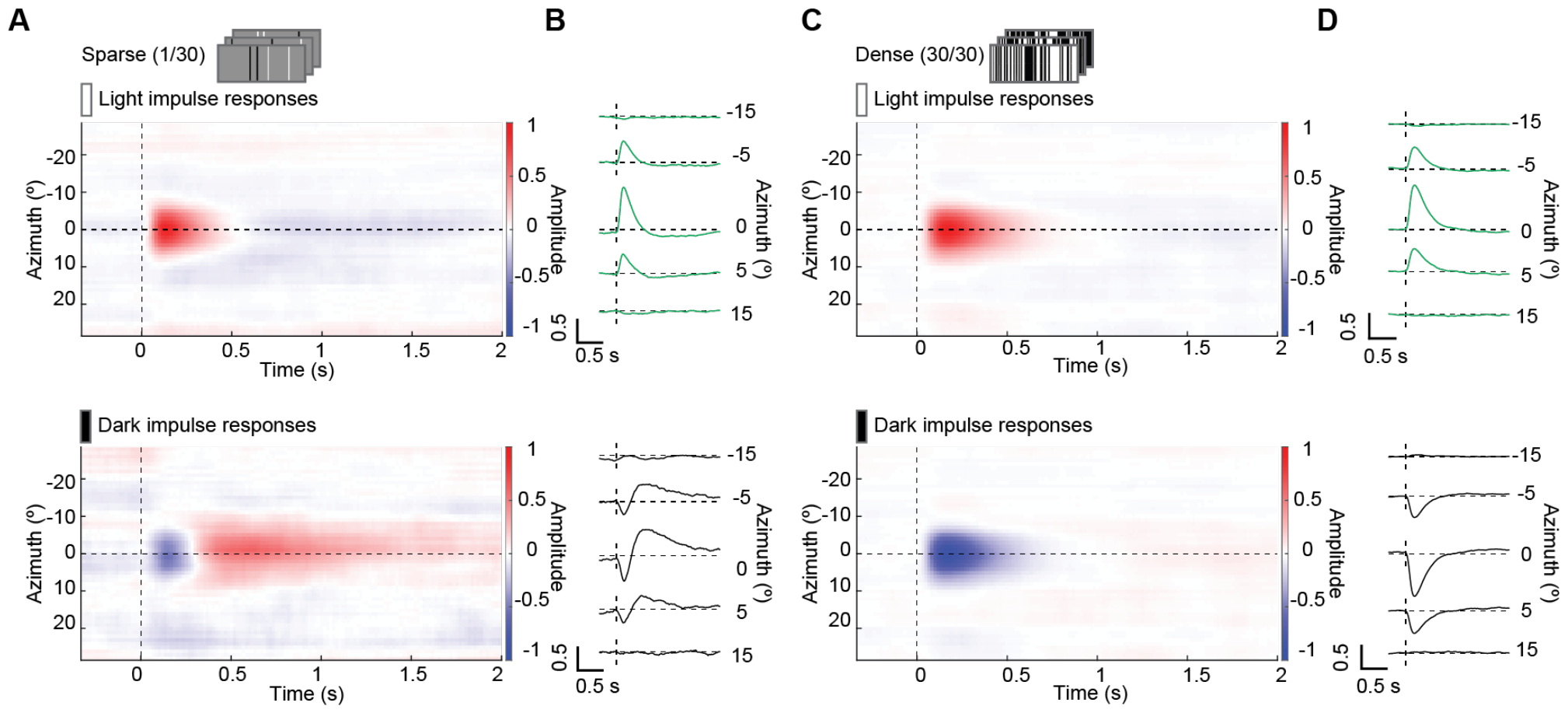
Spatiotemporal impulse responses of Mi1. (Related to Figure 5) (A, C) Average aligned light and dark spatiotemporal impulse responses for sparse (A) and dense (C) stimuli. (N = 14 flies). (B, D) Temporal impulse responses from (A) and (C), slicing the receptive field along the time axis at different locations.

### Sparsity adaptation in Mi1 is localized to its receptive field center

We used full field stimuli to characterize sparsity adaptation in Mi1, which does not account for spatial aspects of response properties. However, neurons upstream to T4 and T5, including Mi1, possess significant antagonistic surrounds ^48,72^. We therefore wanted to understand how sparsity adaptation affects Mi1’s spatial response properties. To measure the spatiotemporal receptive field of Mi1, we employed a one-dimensional ternary-noise stimulus consisting of vertical static bars (each spanning 5°) covering the entire screen (**Fig. 5A**). We varied the sparsity of these stimuli by altering the fill fraction of light/dark frames for each bar. We recorded calcium responses of Mi1 elicited by these stimuli, and subsequently calculated the spatiotemporal RFs using reverse correlation (**Fig. 5B**, see **Methods**). The spatial components of the spatiotemporal RFs are similar in shape and size to previous studies ^48^ in both sparse and dense conditions. To visualize the temporal components of the receptive field at different locations, we plotted receptive field cross-sections at different locations (**Fig. 5C**). The temporal filters at the receptive field center closely match the earlier calcium measurements: they are biphasic under sparse conditions and monophasic under dense conditions. To quantify the impact of sparsity adaptation on spatial response characteristics, we measured the mean receptive field values at different spatiotemporal locations for each fly and compared them between sparse and dense conditions (**Fig. 5D**). The inhibitory surrounds were not significantly different between the two conditions (**Fig. 5d** left). On the other hand, the mean values in the receptive field center from 0.5 to 1 second after the stimulus depended strongly on the stimulus sparsity (**Fig. 5D** right). The receptive field center has a prominent negative lobe under sparse conditions, but is monophasic under dense conditions. Additionally, we calculated the spatiotemporal light and dark impulse responses under sparse (**Fig. S5A, B**) and dense (**Fig. S5C, D**) conditions. These results indicate that impulse responses at the receptive field center align with the previously observed pattern; they differ between the light and dark cases when the stimulus is sparse and under dense conditions, impulse responses are primarily roughly equal and opposite. Stimulus sparsity did not elicit changes in impulse responses in Mi1’s surround receptive field.

**Figure 5.**
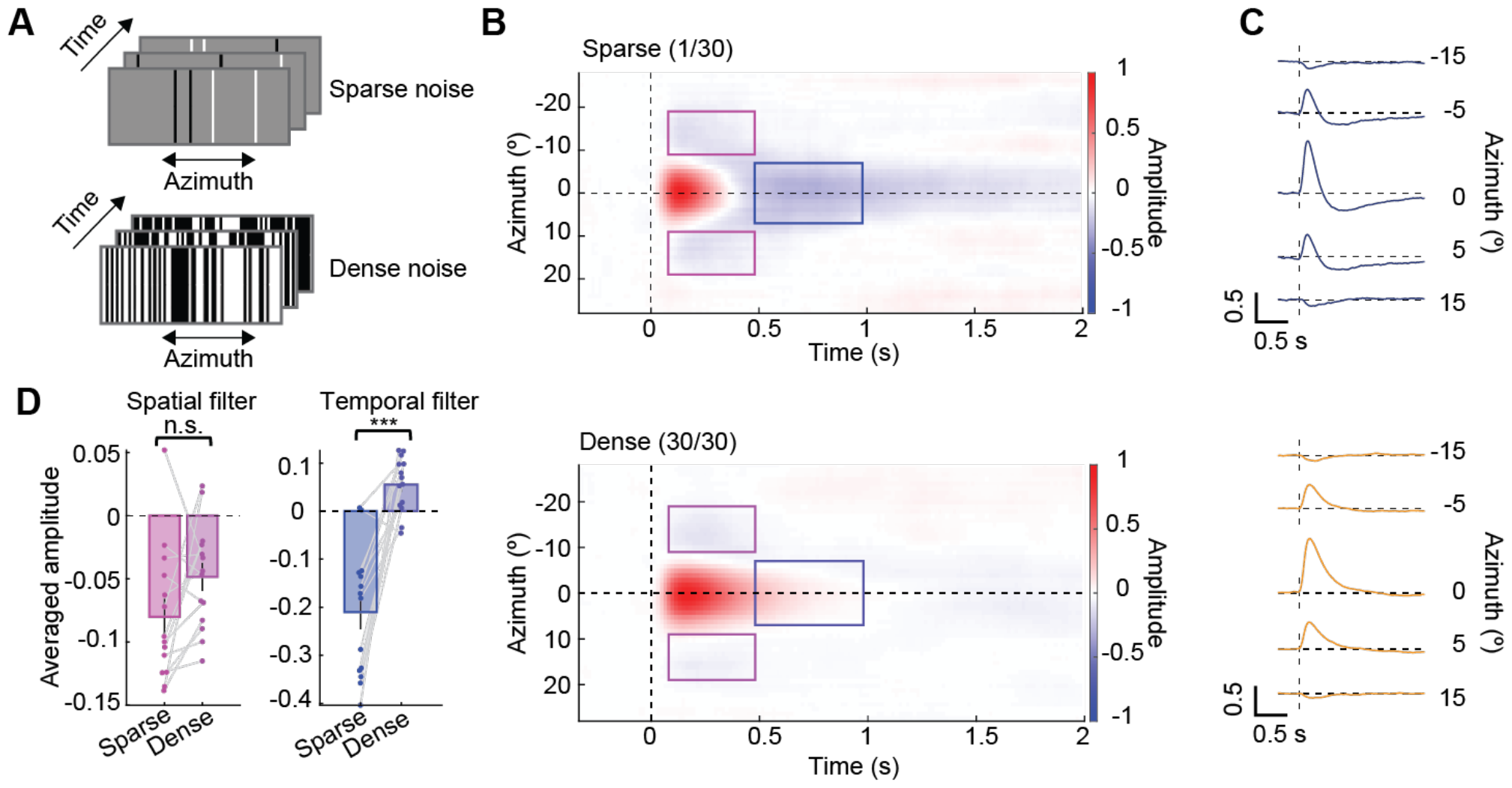
Spatial specificity of sparsity adaptation. (A) Schematic of the sparse and dense one-dimensional ternary-noise stimuli to map the Mi1 spatiotemporal receptive fields. (B) Average aligned spatiotemporal receptive field of Mi1 for sparse (top) and dense (bottom) stimuli. (N = 14 flies). (C) Temporal kernels from (B), slicing the receptive field along the time axis at different locations. (D) Comparison of the spatial and temporal components between sparse and dense conditions through the mean values within the magenta and blue locations in the receptive field. Error bars and shading around mean traces all indicate standard error of the mean. n. s.: not significant (p > .05); *: p < .05; **: p < .01; ***: p < .001; ****: p < .0001 in Wilcoxon signed-rank tests across flies.

**Supplementary Figure 6.**
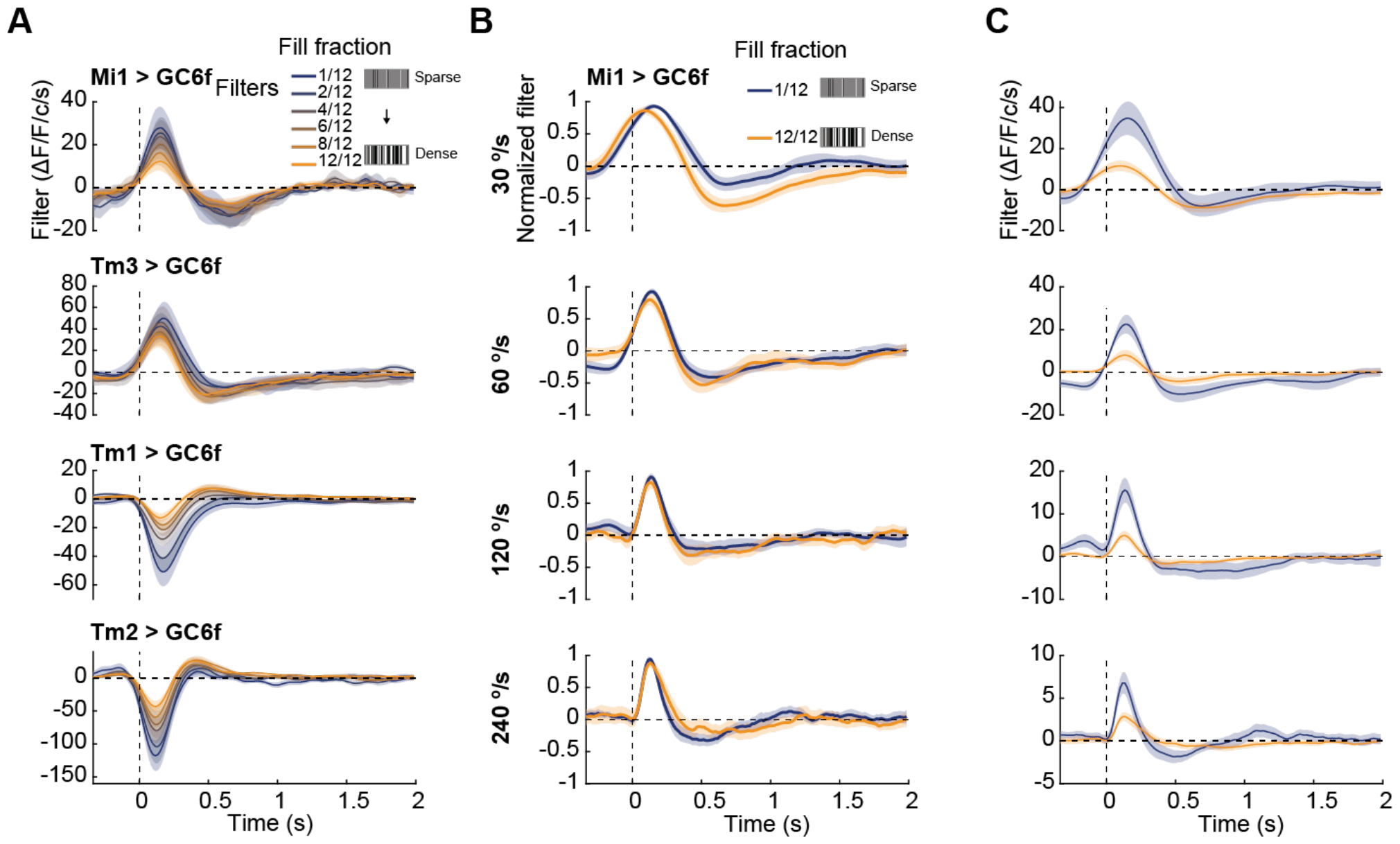
The speed of moving bar stimuli can alter the filter shape change trend. (Related to Figure 6) (A) Linear filters extracted from Mi1, Tm3, Tm1, and Tm2 calcium responses to the moving bar stimuli. Unnormalized version of Fig. 6Ci-Fi. (B) Linear filters extracted from the responses to sparse and dense moving bar stimuli moving at different speeds. All the filters were normalized by the max amplitude. (N = 10 flies). (C) As in (B), except filters were unnormalized. Error bars and shading around mean traces all indicate standard error of the mean.

### Neurons adapt to the sparsity of moving stimuli

To explore sparsity adaptation in a more natural context, we developed a moving bar stimulus in which we could vary the sparsity. In this stimulus, sparse or dense spatial arrays of light/dark or gray bars traverse the screen, creating spatiotemporally sparse or dense inputs to neurons that receive signals from limited regions on the screen (**Fig. 6A**). Each bar had a width of 5° and all moved at a speed of 60°/s. Consequently, 12 bars per second passed each spot on the screen. Sparsity was defined as the fill fraction of light/dark bars to the total number of bars. For instance, a fill fraction (sparsity) of 1/12 corresponds to on average 1 light or dark bar passing per second, while a fill fraction of 12/12 indicates a constant presence of light/dark bars passing a specific location on the screen (**Fig. 6B**). This stimulus thus contains persistent visual objects that move across the retina.

**Figure 6.**
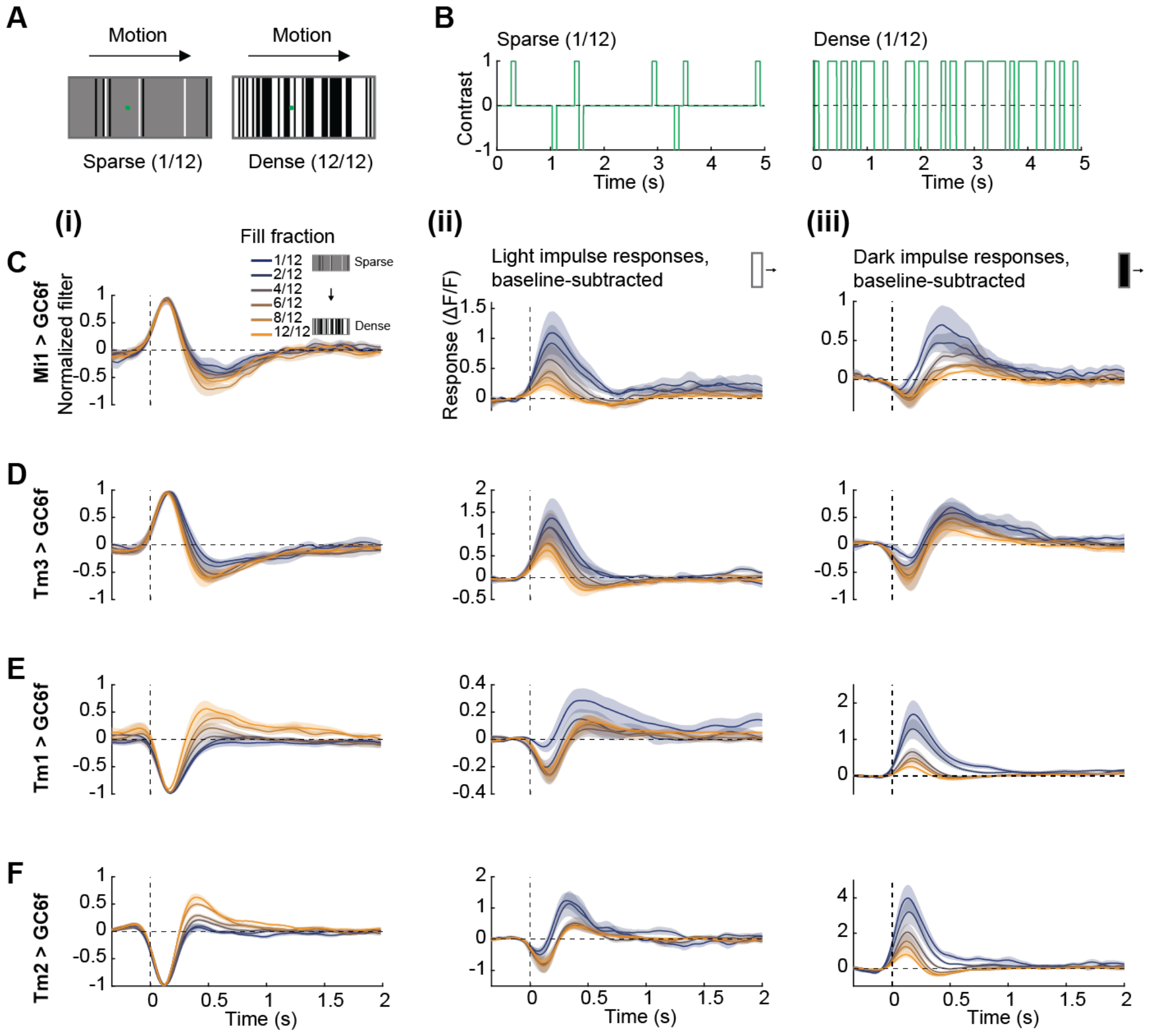
Calcium measurements on the main input neurons to T4 and T5 using moving bar stimuli. (A) Illustration of the sparse and dense moving bar stimuli. Stimuli are stochastic in space and move with a constant velocity. (B) Example time traces of visual contrast at single points on the screen (shown as green spot in A). (C) (i) Linear filters extracted to different fill fraction (sparsity) stimuli from the Tm3 calcium responses to the moving bar stimuli. All the filters were normalized by the max amplitude. (ii) Baseline subtracted light bar responses calculated from the calcium responses of Mi1. (iii) Baseline subtracted dark bar responses calculated from the calcium responses of Mi1. (N = 8 flies). (D) As in (C), except results were calculated from Tm3 calcium responses. (N = 5 flies). (E) As in (C), except results were calculated from Tm1 calcium responses. (N = 8 flies). (F) As in (C), except results were calculated from Tm2 calcium responses. (N = 10 flies). Error bars and shading around mean traces all indicate standard error of the mean.

We measured responses in Mi1 to this new stimulus. As in previous analyses, we computed both filters and impulse responses based on the calcium responses, triggering in time based on when each bar swept through a cell’s receptive field (**Fig. 6C**). The impulse responses resembled those obtained using the earlier, full field ternary noise stimulus (**Fig. 6Cii, iii**). Under sparse conditions, light and dark impulse responses looked very different. Light bars result in transient activity that returns quickly to baseline. The response to dark bars is initially negative with a delayed positive peak, much like earlier sparse impulse responses. In contrast, under dense conditions, the light bar response is a sign-inverted version of the dark bar response. Thus, sparsity adaptation of impulse responses to moving bars bears striking similarity to sparsity adaptation to full screen stimuli. When filters were computed, their shape changed with changing fill fraction (**Fig. 6Ci, S6A**), but interestingly the trend is reversed compared to earlier measurements, so that stimuli with higher fill fraction elicited increasingly biphasic responses. We hypothesized that this difference was due to interactions between the receptive field center and surround, which matters in the case of moving bars but not when the entire screen was uniform, as in our original stimuli (**Figs. 1-5**). To test our hypothesis and elucidate this difference, we varied the velocity of the moving bars from 30°/s to 240°/s under sparse and dense conditions and extracted the filters in each case (**Fig. S6B, C**). At slow speeds, the denser stimuli yielded more biphasic filters, but at high speeds, where a passing bar activates center and surround more simultaneously — and thus more like our original stimulus — the sparser stimuli yielded more biphasic filters. Thus, the speed of bars through the receptive field combines with changes in the receptive field structure with changing sparsity to generate the observed filter shapes.

Since we wanted to examine the impact of sparsity adaptation on motion circuits, we also examined sparsity adaptation in other neurons upstream of the motion detectors T4 and T5 using the moving bar stimulus. The neuron Tm3 showed similar results to Mi1 in its filter and impulse responses changes with sparsity (**Fig. 6D**). In the OFF pathway, both Tm1 and Tm2 also showed robust sparsity adaptation to the moving bar stimulus (**Fig. 6E, F**). Their filters varied with changes in sparsity, while light and dark bar responses differed characteristically in shape under sparse conditions. Overall, moving stimuli induce substantial sparsity adaptation in neurons upstream of motion detectors T4 and T5. This suggests that stimulus sparsity may also strongly affect the downstream computation of motion in T4 and T5.

**Supplementary Figure 7.**
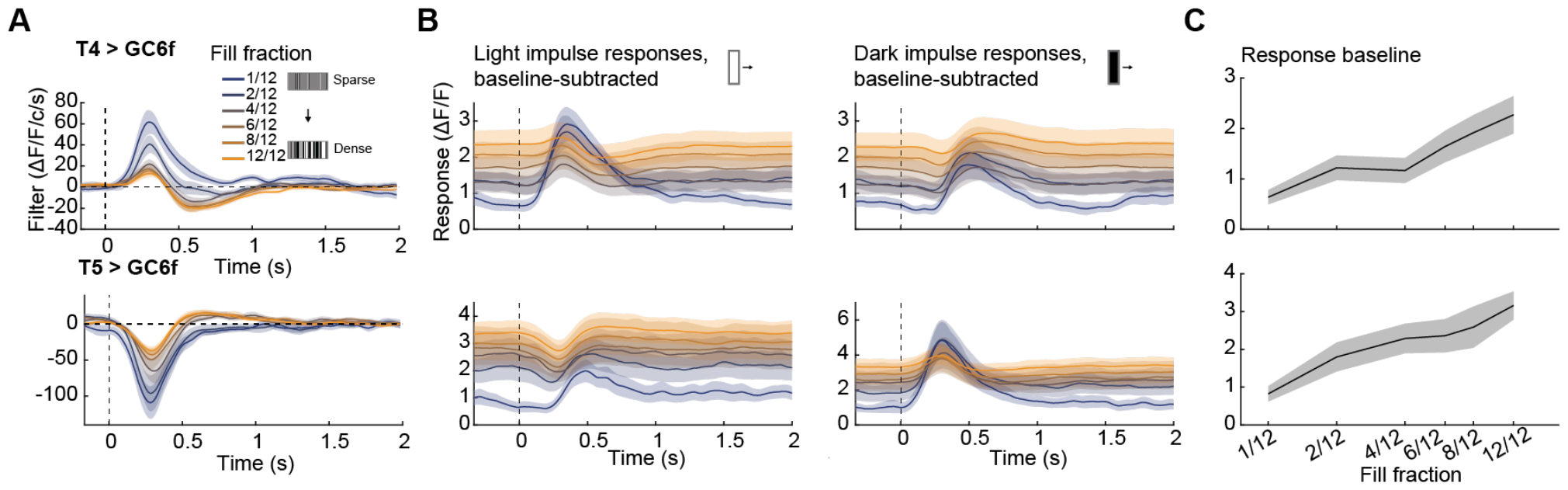
Additional measurements of T4 and T5 responses to moving bar stimuli with different fill fractions. (Related to Figure 7) (A) Linear filters extracted from T4 and T5 calcium responses to moving bar stimuli with different fill fractions, unnormalized version of Fig. 7C, E. (B-C) Light and dark bar responses (B) and baseline responses (C) calculated from the calcium responses of T4 and T5 to moving bar stimuli with varying fill fraction. Error bars and shading around mean traces all indicate standard error of the mean.

### Differential tuning to light and dark isolated moving bars

Motion detection depends critically on the timing upstream neurons ^73,74^. For instance, velocity tuning in T4 is altered when genetic techniques manipulate timing in Mi1 or Tm3 ^75^. Since light and dark impulse responses to sparse stimuli have different shapes in time, we reasoned that sparsity adaptation should alter velocity tuning in T4 and T5 neurons. In particular, we hypothesized that isolated (sparse) moving light and moving dark bars should elicit different speed tuning curves in T4 and T5, due to the different timing in upstream neurons to these stimuli. We therefore measured the responses of T4 and T5 to isolated light and dark bars moving at a variety of speeds in the cells’ preferred direction. The response time traces revealed that T4 responds to light bars (its preferred contrast, PC), but also to dark ones (its non-preferred contrast, NC), while T5 responds to dark bars (its PC), but also to light ones (its NC) (**Fig. 7A**), in agreement a prior report ^76^. For both T4 and T5, the peak time in responses to NC stimuli is delayed compared to the peak time to PC stimuli, reminiscent to the responses measured in neurons upstream of T4 and T5 (**Fig. 6**). The peak values for each curve were used to generate velocity tuning curves (**Fig. 7B, left**). In those curves, the maximum responses to PC stimuli are larger than to NC for both T4 and T5 neurons. However, the tuning to PC and NC differed also in their shape, as measured by the curves’ center of mass (**Fig. 7B, right**), showing that NC responses tend to be tuned to slower velocities. Many neurons upstream of T4 and T5 have altered dynamics (**Fig. 6**), but it seems likely that the large, delayed responses to NC stimuli in upstream neurons result in a relative shift to larger T4 and T5 responses at slow velocities. These results show how the distinct dynamics of light and dark impulse responses to sparse stimuli influence downstream motion computations.

**Figure 7.**
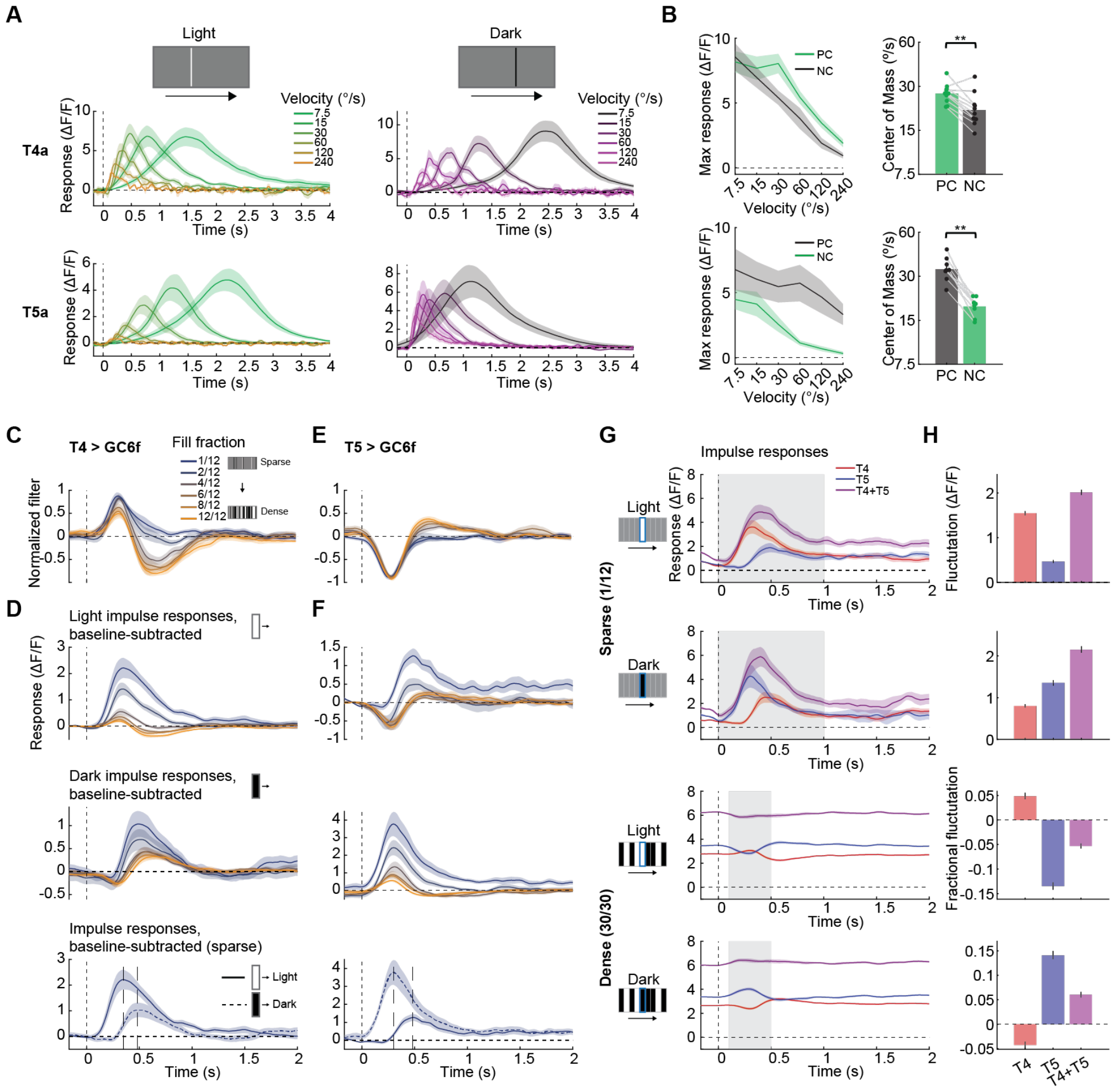
Sparsity adaptation in directional neurons T4 and T5. (A) Aligned calcium responses of T4a and T5a neurons to isolated bars moving in their preferred direction with different velocities. (T4 > GC6f: N = 10 flies; T5 > GC6f: N = 9 flies). (B) Left: T4 and T5 velocity tuning curves to single moving bar were computed from peak values for each curve in (A). Right: The preferred-contrast (PC) and non-preferred-contrast (NC) tuning curves were compared through their center of mass, a weighted average of the tuning curve shown in the left (see **Methods**). (T4 > GC6f: N = 11 flies; T5 > GC6f: N = 9 flies). (C) Linear filters extracted from T4 calcium responses to moving bar stimuli with different fill fractions. All the filters were normalized by the max amplitude. Only responses to preferred directions were plotted here. (N = 11 flies). (D) Baseline-subtracted light (top) and dark (middle) bar responses calculated from the calcium responses of T4 to moving bar stimuli with different fill fractions; baseline-subtracted light and dark bar responses to the sparse stimuli overlaid (bottom). (E) As in (C), except results were calculated from T5 calcium responses. (F) As in (D), except results were calculated from T5 calcium responses. (N = 12 flies). (G) Sparse and dense impulse responses of T4 and T5, plus their simulated sum. (H) The fluctuations of T4 and T5 calcium responses and their sum were compared by computing the mean of the shaded area in (G). In the dense case, the fluctuations were calculated as fractional changes relative to baseline. Error bars and shading around mean traces all indicate standard error of the mean. n. s.: not significant (p > .05); *: p < .05; **: p < .01; ***: p < .001; ****: p < .0001 in Wilcoxon signed-rank tests across flies.

### Sparsity adaptation in T4 and T5 matches properties in upstream neurons

We next sought to characterize T4 and T5 responses to moving stimuli with a range of different sparsities. We measured the calcium responses of T4 and T5 using the earlier moving bar stimulus (**Fig. 6**) with varying fill fractions, which alter the stimulus sparsity (**Fig. 7C**). We analyzed responses to movement in each cell’s preferred direction (see **Methods**). We first extracted the filters describing the response to both light and dark bars at each fill fraction. In T4 and T5 cells, the baseline calcium response relates to the strength of the presented motion signal ^50^. As the fill fraction increased, the baseline response of T4 increased, as expected from the stronger motion signal in the stimulus (**Fig. S7B-C**). In these experiments, filters and impulse responses measure deviations from this baseline, direction-selective increase in calcium. The shape of the T4 filters changed substantially as the fill fraction increased (**Fig. 7cC, S7A**), moving from monophasic to biphasic in a pattern that matches Mi1 and Tm3 filters to the same stimulus at the same speed (**Fig. 6C, D**). Similarly, the light and dark impulses change with sparsity in the same way as those in Mi1 and Tm3 change (**Fig. 7D**, compared to **Fig. 6C, D**). In the case of sparse stimuli, the light bars elicit immediate activation of T4, while dark bars initially reduce activation before activating with a delay. T5 demonstrates similar sparsity adaptation properties, albeit with inverted polarity (**Fig. 7E, F, S7**), which closely resemble responses in Tm1 and Tm2 to identical stimuli (**Fig. 6E, F**). The resemblance between the responses of neurons T4 and T5 and upstream neurons suggest that with these stimuli, T4 and T5 inherit properties from excitatory inputs.

### Sparsity adaptation enhances motion responses to sparse stimuli while suppressing fluctuations to dense stimuli

Last, since sparsity adaptation is prominent in directional circuits, we wondered how it might affect motion detection overall. T4 and T5 neurons provide inputs to many downstream neurons, which often add together their signals and integrate them over space. For instance, horizontal and vertical system cells add together inputs from T4 and T5 and control body and head turning ^50,77,78^. We therefore asked how sparsity adaptation affects the *summed* motion signals from T4 and T5. We approximated the relative responses of potential downstream neurons by simply summing the measured responses of T4 and T5 (**Fig. 7G**). In the sparse case, both T4 and T5 are activated by bars of their preferred contrast, but sparsity adaptation results in activation by bars of their non-preferred contrast, with a short delay. When T4 and T5 signals are summed, the activations add, creating a strong downstream signal. Here, the sparse stimuli result in both T4 and T5 responding to both light and dark stimuli, enhancing signals relative to a case in which each responded to only to bars of their preferred contrast. On the other hand, in the dense case, the light bar response in T4 is approximately equal and opposite to the light bar response in T5, and the responses of T4 and T5 are similarly opposite for dark bars in dense stimuli. Therefore, when T4 and T5 are summed in the dense case, their responses to light bars mostly cancel one another out, and the same holds for dark bars. Since T4 and T5 are constitutively activated by dense stimuli in their preferred direction, this cancellation acts to create only small fluctuations in downstream signals, rather than the larger fluctuations in T4 and T5 alone. This result can be quantified by measuring the fluctuation amplitude in T4, in T5, and in their sum (**Fig. 7H**). Thus, adaptation to stimulus sparsity enhances motion signals to isolated light and dark stimuli by causing more of the visual circuit to respond to those stimuli. And in dense stimuli, the different pattern of responses creates fluctuations in the ON and OFF motion pathway that mostly cancel out when summed.

## Discussion

In summary, we found substantial spatial variability in the sparsity of natural scenes. An early visual neuron in fly’s motion visual system changes its response timecourse dramatically with visual stimuli of different sparsities (**Fig. 1**). This adaptation is distinct from contrast adaptation and depends on the stimulus sparsity, rather than the infrequency of changes in the stimulus (**Fig. 2**). A biophysically plausible linear-nonlinear-linear (LNL) model reproduces the observed sparsity adaptation phenomenology, due largely to a saturating nonlinearity, suggesting a general mechanism for such phenomena (**Fig. 3**). But optogenetic experiments show that this model does not map neatly onto the voltage-to-calcium transformation in early visual neurons (**Fig. 3**). Indeed, sparsity adaptation was widespread in the visual system, including in both ON and OFF pathways (**Fig. 4**). The adaptation affected the receptive field center in Mi1 but not its surround (**Fig. 5**). Moving sparse and dense stimuli could also elicit changes in processing across the visual system upstream of motion detectors (**Fig. 6**). Furthermore, the motion detectors T4 and T5 showed similar processing changes, including a predicted difference in the velocity tuning of both T4 and T5 with stimulus polarity. Overall, sparse light and dark stimuli elicit positive responses in both ON and OFF pathway circuits, but dense stimuli elicit responses with opposite signs in the two pathways, so that one pathway is active when the other is suppressed (**Fig. 7**). Thus, overall, dense light and dark stimuli lead to oppositely signed responses in the ON and OFF pathways, while sparse light and dark stimuli recruit responses both ON and OFF circuitry with can sum constructively, independent of the stimulus polarity.

### Relationship between sparsity and other statistical properties

Prior work has investigated visual adaptation to stimuli by examining how the different moments of the intensity distribution affect processing properties. Notably, the stimulus mean (first moment) has strong effects on spatial and temporal receptive fields ^17,19,30,31^, while the stimulus variance (second moment) strongly affects response amplitudes ^14,15,20,21,56^, as well as dynamics ^14,21,34-36^. Other studies have shown that vertebrate retinas adapt to spatiotemporal correlations in intensity ^37^. Studies in both salamander retinal ganglion cells ^23^ and cat lateral geniculate nucleus neurons ^22^ revealed strong adaptation to changes in first and second order statistics, but did not observe strong adaptation to the higher moments of the luminance distribution, specifically to its skewness (third moment) or kurtosis (fourth moment). Other studies have shown strong adaptation to phase correlations ^24,25^, which correspond to spatial or temporal structures, like edges, in the stimulus. On the other hand, the present study demonstrated that neurons in fly motion detection circuits exhibit strong adaptation to the statistical property of sparsity (**Figs. 1, 4, 5, 6, 7**). Notably, changes in sparsity affect the stimulus variance, kurtosis, and other, higher even-ordered moments of the distribution, and notably alter the spatial and temporal structure of the stimulus, though not the shape of its power spectrum. The effects of sparsity are distinct from contrast adaptation (**Fig. 2**), so the visual system must be adapting to a combination of (even) higher order statistics of the stimulus captured by its sparsity. Since our implementation of changing sparsity relates to only even moments of the distribution, the effects we see are distinct from prior studies of third-order correlations, which can signify motion and are perceived by both flies and humans as motion signals ^79-82^.

The phenomenological model we introduced could reproduce our qualitative results in Mi1 and must generate sensitivity to signal sparsity, among other features. It is reminiscent of prior modeling studies that introduced saturating nonlinearities that improved the accuracy of motion responses to natural scenes ^21,83^. Thus, it also seems plausible that the adaptation to stimulus sparsity is consistent with improving motion detection under natural conditions. In natural scenes, many statistical features vary over space, so adapting to sparsity, in addition to changing contrasts, may improve the fidelity of motion detection.

Compared to considering higher moments of intensity distributions or phase correlations, sparsity has the advantage that it can be simply interpreted: sparse scenes are those in which relatively isolated objects stand out against relatively uniform backgrounds, while dense scenes are those in which there are lots of local visual features. Adaptation to sparsity consists of changes in processing related to these two distinct types of visual scene. Interestingly, the principle of sparse-coding suggests that sensory systems may encode stimuli using only a small subset of neurons ^40,42,84^. In contrast, in this study, the adaptation of the visual system to sparse stimuli made the response *less sparse* by recruiting responses in a larger set of neurons.

### Sparsity in ON and OFF pathway segregation

ON and OFF channels, which respond to stimulus increments and decrements, respectively, are prevalent across sensory systems, from vision ^85^ to taste ^86^ to audition ^87^. This split into ON and OFF pathways is thought to confer a metabolic advantages ^88,89^ and to enhance the efficiency of stimulus encoding ^90^. In vision, ON pathways have positive preferred contrasts, since they respond to luminance increments, while OFF pathways have negative preferred contrasts, since they respond to luminance decrements. However, a primary result in this paper is that the boundary between ON and OFF channels depends on the sparsity of the stimulus. In particular, ON- and OFF-motion detectors T4 and T5 can be driven positively by isolated bars of both polarities moving in their preferred direction, even while they respond with opposite signs to light and dark bars in dense stimuli (**Fig. 7**). This is in agreement with a prior report that T4 and T5 respond to bars of their non-preferred contrast ^76^. Importantly, our data show that these properties depend critically on the sparsity of the stimulus (**Fig. 7D, F**) — for instance, in dense stimuli, T4 and T5 neurons are not activated by their non-preferred contrast. Moreover, the responses in T4 and T5 mirror the response properties of cells upstream of T4 and T5, suggesting that they inherit many of these properties and do not require distinct conductances to generate them, as was proposed to explain the earlier observation ^76^ (**Fig. 6**).

Upstream of T4 and T5, we observed a similar blurring of the ON and OFF pathway properties under sparse conditions. In dense conditions, ON (OFF) cells become more active in response to light (dark) flashes and bars and less active in response to dark (light) flashes and bars (**Fig. 1, S1, 4, S4, 6**). In sparse stimulus conditions, ON (OFF) cells respond positively to both light and dark isolated pulses, with the response to the non-preferred contrast delayed relative to the preferred one (**Fig. 1M, 4A-F, S4A-C, 6C-F**). This effect has been observed in a prior study ^55^, in which Mi1 and Tm3 responded positively to both light and dark isolated flashes, with the pattern seen here. Importantly, our data show that this effect occurs on a spectrum from sparse to dense stimuli and the changes in ON and OFF responsivity affect downstream processing. The delayed activation to non-preferred contrasts appears in voltage measurements, upstream of calcium signals (**Fig. 4Biii**), does not depend on the calcium level in a cell (**Fig. 3L, M**), and appears in neurons upstream of inputs to T4 (**Fig. 1M, 4Diii, 6C, D**). Most importantly, our data suggest that these properties also translate into the sparse responses of T4 and T5, conferring on them sensitivity to non-preferred contrasts of objects moving in their preferred directions. A different study observed more bilobed responses to isolated light flashes and used linear filters based on those flash responses to simulate neural responses and compare them to T4 and T5 responses ^91^. Here, we note that the preferred and non-preferred contrast flash responses have very different shapes (**Fig. 1, 2, S2, 4, S4)**. That is, the system is far from its linear regime during these responses. And since the impulse response dynamics depend on the stimulus statistics, it is difficult to use the impulse responses to predict responses to other stimuli with different sparsity.

Blurring of the ON and OFF dichotomy in response properties has been observed in early vertebrate visual systems. For instance, in salamander retina, OFF-type cells can be driven to have ON-type responses by stimuli far from their receptive field ^92^. Similarly, in mouse retina, as the ambient light level changes, many OFF-type retinal ganglion cells acquire responses to increments and many ON-type cells acquire responses to decrements ^33^, both delayed in a manner similar to the non-preferred contrast responses observed here. Thus, stimulus statistics may very generically affect the ON and OFF channel encoding of visual stimuli and alter the properties of cells often typed as exclusively ON or exclusively OFF. Here, we show that the changes in encoding with stimulus sparsity serve to recruit more neurons to respond to stimuli that are infrequent in time or space, a potential mechanism for enhancing responses to sparse stimuli.

### Sparsity adaptation could act as a mechanism for bottom-up saliency enhancement

Sensory systems receive vast amounts of data, and processing this information poses a challenge for a brain. To meet the challenge, brains deploy attention, which selectively concentrates neural resources on specific aspects of sensory input or behavioral tasks ^93,94^, emphasizing certain signals over others ^95^. Attention can be voluntarily deployed (top-down) or generated by the physical salience of a stimulus (bottom-up) ^96,97^. Models of bottom-up attention rely on creating stronger neural responses to some stimuli than to others, thus creating a ‘saliency map’ of stronger versus weaker neural signals ^98-100^. This sort of bottom-up modulation of neural signal strength over time and space has been observed in visual cortex in response to pop-out stimuli ^101-103^ (i.e., stimuli with visual features distinct from surrounding distractors) and abrupt onset stimuli ^104-106^. The sparse stimulus responses we observe recruit both ON and OFF pathway neurons to respond to isolated light or dark moving objects, ultimately increasing the response of the visual system to these objects compared to a hypothetical system that had only ON or only OFF channels responding to light or dark moving objects, respectively. In contrast, the dense stimuli generate opposing responses in ON and OFF channels. Thus, the processing observed here acts to increase the salience of sparsely distributed objects and their motion in visual scenes, perhaps as a distinct form of pop-out response.

Sparsity adaptation is likely to impact many downstream signals and behaviors. Optomotor responses rely on motion signals from T4 and T5 neurons ^50,51^, and sparsity adaptation will tend to enhance motion signals from isolated objects; this will increase optomotor response strength compared to a hypothetical visual system that did not recruit both ON and OFF channels to sparse stimuli. Because the tuning to non-preferred contrast is to slower speeds than for preferred contrast, this effect should be more pronounced at low speeds (**Fig. 7**). Other detectors that lie downstream of T4 and T5, such that those for looming ^107^ and small object avoidance ^108^, should also benefit from enhanced signals in sparse compared to dense scenes. Similarly, small object detectors that are sensitive to ON and OFF signals, like LC11 ^109,110^ would have their signals enhanced by this effect. Overall, by recruiting both ON and OFF channels to encode sparse light or dark stimuli, sparsity adaptation acts to boost neural signals and increase the salience of spatially and temporally infrequent signals.

## Acknowledgements

This study was supported by NIH R01 EY026555 and NS121773. We benefited from helpful conversations with Jonathan Demb, Ryosuke Tanaka, and Baohua Zhou.

## Contributions

Original studies were conceived by TG, CAM, and DAC. CAM obtained and analyzed early data. TG performed experiments, modeling, and data analysis. TG and DAC wrote the paper.

## Methods

### Resource Table

| Reagent or resource | Source | Identifier |
| --- | --- | --- |
| **Chemicals** |  |  |
| All-trans retinal | Sigma Aldrich | PubChem SID: 24899355 |
| **Model organisms** |  |  |
| *D. melanogaster*: Mi1: +; +; R19F01-Gal4 | 46 | BDSC: #48852 |
| w; UASGC6f; + | 52 | BDSC: #42747 |
| w; +; UASGC7b | 57 | BDSC: #79029 |
| +; +/CyO; ArcLD/TM6B | 62 | BDSC: #51057 |
| +; +; iGluSnFR | <sup>111</sup> (Gift from Marc Freeman) | N/A |
| w-; UAS-GACH3.0; + | <sup>70</sup> (Gift from Yulong Li) | BDSC: #86549 |
| +; UAS-ChrimsonR; + | 61 | BDSC: #55136 |
| +; L1(c202a)-Gal4; + | 112 | N/A |
| +; +; 0595-Gal4 | 113 | N/A |
| w; R21A05-AD; R31H09-DBD | 114 | N/A |
| w; R41G07-AD; R74G01-DBD | 68 | BDSC: #86810 |
| w; R28D05-AD; R82F12-DBD | 68 | BDSC: #86811 |
| +; +; R13E12-Gal4 | 47,115 | BDSC: #48569 |
| w, nSyb-phiC31; +; + | 116 | BDSC: #84150 |
| w; R59E08-AD; R42F06-DBD (split T4/T5) | 117 | JRC: SS00324 |
| +; UAS-SPARC-I-GC6f; + | 116 | BDSC: #84138 |
| **Software** |  |  |
| MATLAB | Mathworks | N/A |
| Psychtoolbox 3 | 118-120 | N/A |
| ScanImage 5 | 121 | N/A |
| **Other** |  |  |
| Natural scene images | 44 | N/A |

### Natural scene characterization

To characterize sparsity in natural scenes, we chose natural images from a database ^44^. This dataset contains panoramic images captured in diverse environments, each measuring 360×97.5 degrees and sampled at approximately 2.6 pixels per degree. We chose 44 natural images that excluded indoor and architectural scenes. Prior to calculating the sparsity index, we preprocessed the selected images. This involved cropping each image into small patches using sliding square windows of dimensions 5, 15 or 41 degrees. In each patch, we inserted a black pixel at its center and then computed the local contrast, denoted as *c*(*x, y*), at each location (*x, y*) using the following equation:

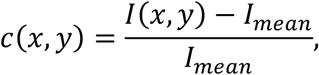

where *I*(*x, y*) is the luminance at each location, and *I*_*mean*_ is the mean luminance of this patch. Sparsity is a metric to describe whether the energy of a signal is concentrated in a small number of significant components or evenly distributed ^122^. Quantitatively, it can be measured using Gini index ^45^, negative entropy ^123^, or by the sparsity index proposed below. The sparsity index for this patch was calculated by dividing the mean absolute contrast by the root mean contrast of the pixels in the patch (this is related to the ratio of the *L*^*1*^ and *L*^*2*^ norms):

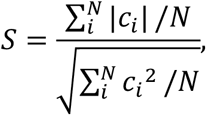

where *c*_*i*_ is the contrast of each element within this patch, and *N* denotes the total number of elements within the patch. The black pixel was added to the patch because this metric, *S*, is otherwise not sensitive to the overall scale of *c*_*i*_. Without the black pixel, small fluctuations about a mean luminance in the sky, for instance, will yield a similar value as the large contrast fluctuations in shrubbery. This metric approaches 1 when there is equal power in all pixels, that is, when all *c*_*i*_ are equally far from 0. It approaches 2(*N* − 1)^1*/*2^*/N* when all pixels are constant and non-zero but one is black (*i*.*e*., its luminance is 0).

### Fly strains and husbandry

Flies were grown in incubators at 25ºC. Non-virgin female flies were used for all experiments. Flies were briefly anesthetized with ice before surgery and imaged between 2 and 7 days post-eclosion. For imaging experiments with optogenetics, flies were dark reared on food containing 10 μM all-trans retinal (ATR) ^124^ for at least 2 days. Genotype details are summarized in **Supplementary Table 1**, and parental lines are given in the **Resource Table**.

**Supplementary Table 1.**
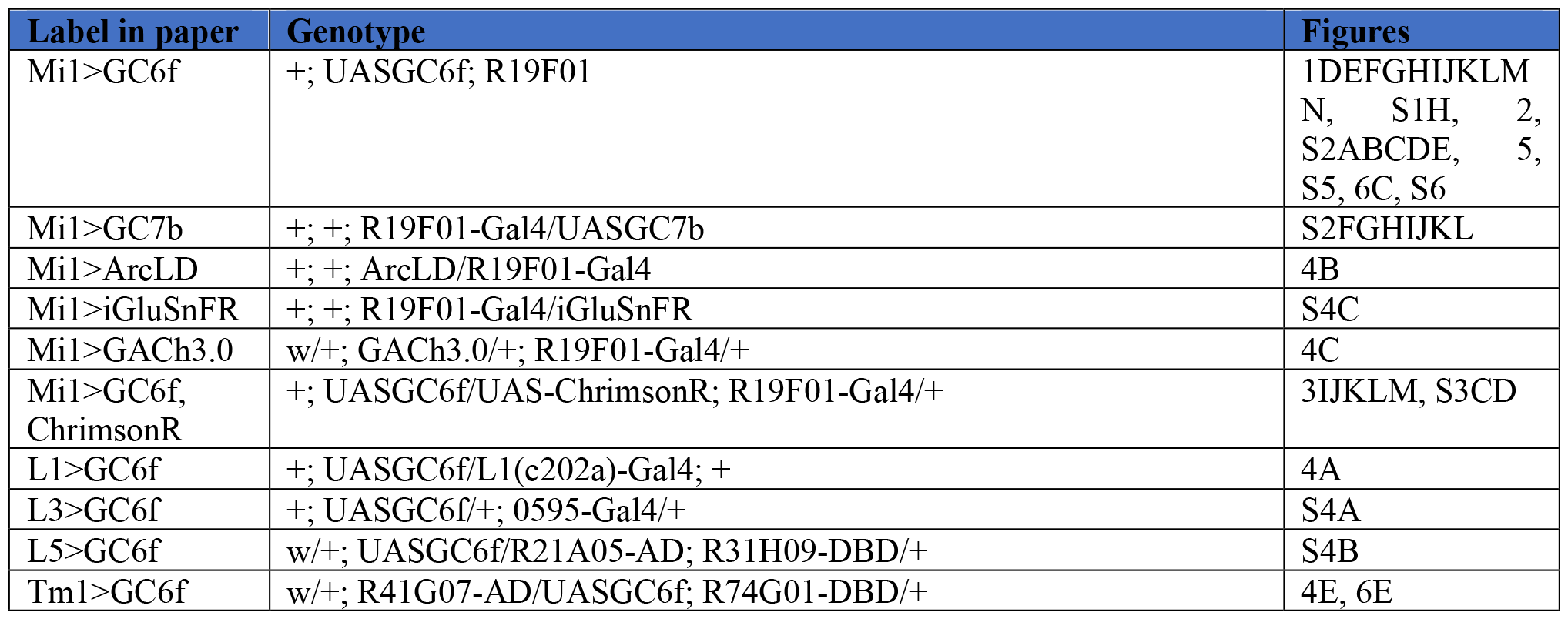
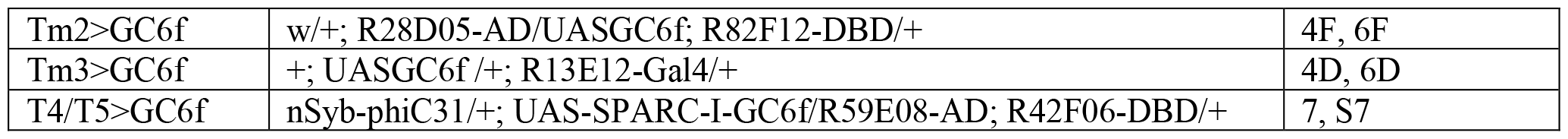
Genotypes of flies used in the experiments.

### Visual stimuli presentation

Visual stimuli were generated using Matlab and Psychtoolbox ^118-120^. Stimuli were presented on a virtual cylinder around the fly using a digital light projector (Texas Instruments, TX, USA) that refreshed the screen at 180 frames per second ^125^. To prevent green light bleeding into photomultiplier tubes (PMT) used for microscopy, the projector output was filtered with two 565/24 nm filters in series (Semrock, NY, USA), while 512/25 and 514/30 filters (Semrock, NY, USA) were used as emission filters in front of the PMT. The mean intensity of stimuli was ∼80 cd/m^2^. Throughout, the gray background was equal to the projector’s half-brightest luminance, and unless specified otherwise, all visual stimuli were presented with this gray background. A compilation of the visual stimuli used in the experiments is provided in **Supplementary Table 2**.

**Supplementary Table 2.** Stimuli used in the experiments and probes used to identify and select regions of interest for analysis.

| Stimulus | Description (duration) | Figures |
| --- | --- | --- |
| Full field ternary noise sparsity sweeping stimulus | Ternary full-field flickering stimulus, updated stochastically at 30 Hz. It consists of different epochs with varying sparsity, determined by the ratio of light/dark flash frames to total frames (fill fraction). In the sparsest epoch, the fill fraction was set to 1/30, indicating an approximate occurrence of one light/dark flash frame per second, with the stimulus updating rate of 30 Hz. In the dense epoch, the fill fraction was 30/30, signifying the absence of gray frames and the exclusive presence of light and dark flash frames. The fill fractions used in this stimulus were 1/30, 2/30, 4/30, 8/30, 15/30 and 30/30. When the screen was not gray, the light and dark screens appeared with equal probability. Each epoch lasted 30 s and epochs were interleaved by 2 s periods of uniform gray. | 1FGHIJKLMN, S1BDE, 2F, S2FGHIJKL, 3, S3, 4, S4 |
| Full field uniform noise sparsity sweeping stimulus | Similar to ternary noise stimulus, with the only difference being the substitution of the light/dark frame by a frame in which contrast values are drawn from a uniform distribution with contrasts ranging from -1 (dark) to 1 (light). | S1FGH |
| Full field ternary noise stimulus with contrast sweeping | Similar to ternary noise stimulus. This stimulus had only fill fractions of 1/30 and 30/30, but the light/dark frames had contrasts of 0.1, 0.2, 0.4, 0.8 and 1. | 2ABCDE, S2ABCDE |
| Full field dense noise stimulus with transition frequency sweep | Modified from ternary noise stimulus by interpolating the gray frames using last light/dark frame contrast to obtain 30/30 fill fraction but with changes in contrast occurring at different frequencies, ranging from 1 Hz to 30 Hz (fill fractions 1/30 to 30/30). | 2GHI |
| Sparse and dense one-dimensional ternary-noise stimulus | Similar to the ternary noise stimulus above, except the screen was covered by vertical bars, each spanning 5° azimuthally. Each bar was updated stochastically at 30 Hz using ternary flicker, independently from other bars. The sparsity was adjusted by altering the fill fraction of light/dark frames for each bar, with the sparse epoch as 1/30 and dense epoch as 30/30. | 5, S5 |
| Moving bar stimulus with | Arrays of light/dark or gray bars traverse the screen, with 5°-wide bars moving at 60°/s. The sparsity was adjusted by the ratio of light/dark | 6, S6A, 7CDEFGH, |
| sparsity sweeping | bars to the total bars passing a location per second. Sparsest: 1/12 (about 1 bar/s); densest: 12/12 (continuous bars). When not gray, bars were equally likely to be light or dark. | S7ABC |
| Sparse and dense moving bar stimulus with velocity sweeping | Same as moving bar stimulus above, but only the 1/12 and 12/12 fill fraction epochs were kept, and their speed was swept over 30, 60, 120 °/s to 240 °/s in **Fig. S6bc**. | S6BC |
| Single moving bar stimulus with velocity sweeping | A single moving bar traversed the screen at speeds of 7.5, 15, 30, 60, 120 and 240 °/s. | 7AB |
| **Selection Probes** |  |  |
| Full-field flashes | Alternating full-field light or dark with a period of 2 seconds. Used to select ROIs of non-direction-selective cell types. | 1, S1H, 2, S2, 3IJKLM, S3CD, 4, S4 |
| Full contrast wide moving bar | A light, 10°-wide bar moved in all four cardinal directions at ~30°/s on a dark background for ON-center receptive fields (RFs) cells (like Mi1 Tm3, L5); dark moving bar on a light background for OFF-center RFs cells (L1, L3, Tm1, Tm2). Used to locate receptive fields. | 1, S1H, 2, S2, 3IJKLM, S3CD, 4, S4 |
| Full contrast fast moving bar | Same as full contrast wide moving bar, but 5° wide and moving at a speed of 60 °/s for extracting the receptive field locations on the screen for different ROIs. | 5, S5, 6, S6 |
| Moving square wave grating | Full-contrast square waves with 30° periods moved in all four cardinal directions at 30°/s for selecting active T4/T5 ROIs. | 7, S7 |
| Moving edges | Light/dark edges moved left or right on a dark/light background at 30°/s for selecting T4/T5 ROIs. | 7, S7 |

### Two-photon imaging

Neural activity was recorded with two-photon scanning fluorescence microscopy. Flies were immobilized in a metal holder using UV cured epoxy, with subsequent surgical removal of the cuticle and fat tissue to expose the brain’s optic lobe. Recordings were consistently conducted on the right hemisphere of the brain, with epoxy used to immobilize the mouthparts and minimize brain movement. Oxygenated sugar-saline solution was used to cover the exposed brain ^126^. The metal holder was positioned above panoramic screens, and imaging was conducted using a two-photon microscope (Scientifica, Clarksburg, NJ). Photomultiplier tube (PMT) input was filtered using a series of filters, including 512/25 and 514/30 (center/FWHM) filters (Semrock, Rochester, NY, USA), which blocked the light used for the visual stimulus. Excitation light at 930 nm was delivered by a femtosecond Ti-sapphire laser (Mai Tai; Spectra-Physics), with typical power less than ∼30 mW at the sample. Images were acquired at a rate of ∼13 Hz using ScanImage ^121^ software and were subsequently motion-corrected offline. Custom Matlab code was employed to process and analyze the data.

### Optogenetic activation during imaging

ChrimsonR optogenetic activation was conducted under the two-photon microscope (**Fig. 3I-M, S3C, D**) using a Thorlabs 690 nm laser diode (Thorlabs, HL6738MG) shone through the imaging objective onto the brain. The laser power at the sample was measured at ∼ 1 μW/mm^2^.

### Imaging data analysis

Prior to ROI (regions of interest) extraction for neurons, we manually selected regions in time-averaged fluorescent images that contained neurons in the desired layers. Then, a watershed algorithm ^127^ was used to extract ROIs from these regions. To eliminate remaining stimulus bleed through, we performed pixel-wise background subtraction for each ROI. Here, a background pixel is defined as any pixel not included in the extracted ROI mask. We subtracted the average pixel intensity of every background pixel within 6 pixels away in the y coordinate from each pixel within every ROI. (This background subtraction is especially important when correlating responses with stimulus intensity.) The fluorescent time trace of each ROI was computed by averaging pixel values within the ROI. Then, to convert the fluorescent time traces into the unit of Δ*F/F*, as described previously ^128^, we calculated the baseline fluorescence *F*_0_(*t*) by fitting an exponential to the time trace for each ROI within each interleave epoch. Then, we subtracted this fitted exponential from the original fluorescent time traces specific to each ROI. The resulting difference ( Δ*F*) was then divided by the same fitted exponential, yielding Δ*F/F* time traces, Δ*F/F* = (*F*(*t*) – *F*_0_(*t*))*/F*_0_(*t*).

Following the extraction of ROIs and calculation of Δ*F/F*, responsive ROIs were selected based on their responses to probe stimuli presented before and after each recording session. For non-direction selective neurons exposed to full-field flicker stimuli (**Fig. 1G-N, S1H, 2, S2, 3I-M, S3C, D, 4** and **S4**), as previously described ^21^, ROIs were selected based on their responses to both full-field flashes and bars moving in all four cardinal directions. In the full-field flash selection process, ROIs were selected based on the mean differential response, μ, calculated as the mean response during full-field light periods minus the mean response during full-field dark periods (which alternated with a 2-second period). For neurons with ON-center receptive fields (RFs), we selected the 30% of ROIs with the most positive values of μ within each fly. Conversely, for cells with OFF-center receptive fields, we selected the 30% of ROIs characterized by the most negative values of μ. Additionally, following the full-field flash selection process, we further refined ROI selection based on their responses to bars sweeping across the field of view in all four cardinal directions. ROIs were selected if their responses to at least two directions of the moving bar surpassed a predefined threshold level. This step guaranteed that selected ROIs represented neurons with receptive fields on the screens.

For non-direction selective neurons exposed to moving bar stimuli (**Fig. 6 and S6**) or the sparse and dense receptive field mapping (**Fig. 5 and S5**) stimuli, as described previously ^108^, ROIs were selected based on their responses to a probe stimuli that contains light/dark bars moving horizontally. This probe stimulus was presented multiple times before and after each recording. Pearson correlations between every pair of responses were calculated to assess the consistency of their responses to probe stimuli. ROIs with an average correlation below thresholds between 0.2 and 0.3 were subsequently excluded. In addition, the horizontal location of each neuron’s RF on the screen was calculated based on the response to the probe. This information was used to align the stimuli across ROIs that responded to different regions of the screen.

For direction selective neurons T4 and T5 (**Fig. 7 and S7**), ROIs were selected based on responses to a probe stimulus consisting of square waves moving in all four cardinal directions, and light and dark edges moving horizontally. Following a previously described procedure ^128,129^, moving square waves were presented multiple times and the consistency (Pearson correlation > 0.3) of ROIs’ responses were used to select responsive ROIs. Light and dark edges moving front-to-back and back-to-front were used to construct direction selectivity indices (DSIs) and edge polarity selectivity indices (ESIs). ROIs with a DSI that met threshold criteria were selected as progressive (DSI > 0.3) or regressive (DSI < -0.3) sensitive neurons; similarly, ROIs with an ESI met threshold criteria were identified as T4 (ON-edge selective, ESI > 0.3) or T5 (OFF-edge selective, ESI < - 0.3). Additionally, we determined the azimuthal location of each ROI’s receptive field based on its responses to these moving edges; this information was used for kernel analysis or aligning response time traces.

### Computing the temporal filter and light/dark impulse responses

For neurons exposed to full field ternary noise sparsity sweeping stimulus, linear filters were extracted for each stimulus epoch with different signal sparsity. We applied ordinary least-squares (OLS) regression to calculate the filter as the weighting of the stimulus that predicted the response with the least squared error. In this approach, we used *N* pairs of stimulus-vectors and scalar responses to solve the equation *Sk* = *r*, where *r* represents a vector over *N* scalar responses, and *S* is a matrix of stimulus values at the time points preceding each response:

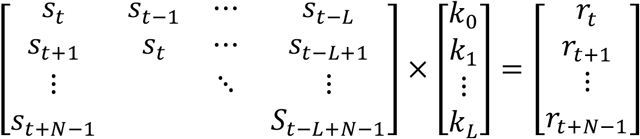

Here, *s*_*r*_ and *r*_*t*_ denote the stimulus and response values at a specific time point. The filter length was set to *L +* 1, and each value *k*_*i*_ of the filter corresponds to the weight with a time lag of *i*.

For neurons exposed to moving bars stimuli, linear filters were extracted for each stimulus epoch with different signal sparsity by calculating the stimulus-response reverse correlation ^54^:

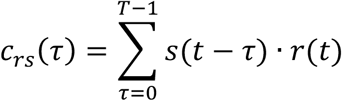

where *s* and *r* represent the stimulus and response vector, respectively, *T* is the length of the kernel to be computed. Then the resulting reverse correlation function was normalized to get the linear filters *k*:

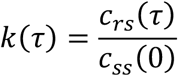

where *c*_*ss*_ and *c*_*rr*_ represent the autocorrelations of stimulus and response vector, respectively.

Light and dark impulse responses were calculated by averaging the responses after a light or dark flash. Specifically, the responses in the time window after each light/dark flash were extracted, and the resulting responses were then averaged to provide an estimate of a neuron’s light/dark impulse response:

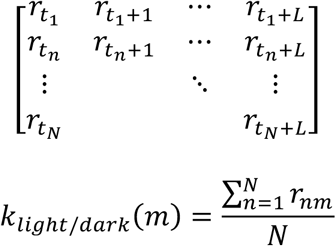

The times *t*_*n*_ were chosen to match when light or dark frames were present on the screen. Note that these impulse responses are closely related to the reverse-correlation filters, but are triggered off of only one type of contrast and do not weight by the contrast polarity.

### Spatiotemporal receptive fields

Spatiotemporal receptive fields were calculated by calculating the stimulus-response reverse correlation:

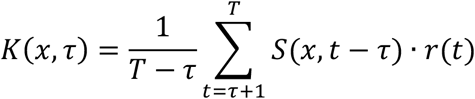

where *S* is the stimulus at each location *x* relative to the ROI receptive field (located based on responses to the moving bar probe stimulus), *r* is the response vector and *T* is the duration of recording.

To compute the spatiotemporal RFs separately for light impulses and dark impulses, at each spatial location, we calculated the light and dark impulse responses separately and subtracted their baselines to generate spatiotemporal light and dark impulse responses.

For display and comparison purposes, spatiotemporal receptive fields from each ROI within flies were averaged to generate each fly’s result. These averages were then normalized for each fly by dividing the maximum amplitude. The results were then averaged over flies.

### Assessing the degree of sparsity adaptation

To evaluate the degree of sparsity adaptation, we introduced a metric called the shape index to quantify filter shape (**Fig. 1H, 2c, S2G, and 3C**). The absolute, unsigned areas of the filter’s first and second lobe were calculated as A and B. The shape index was defined as the ratio of their difference to their sum: (A − B)/(A + B).

### Tuning curves and center of mass

To compute the single moving bar tuning curves for T4 and T5 (**Fig. 7A, B**), we recorded responses to a stimulus consisting of an isolated light/dark bar, 5º wide, moving across the screen at various speeds. For each speed, we aligned the response time traces over all the ROIs based on their computed RF locations on the screen, found from a moving edge probe stimulus. We then averaged the ROIs within flies to generate each fly’s response time trace. Then, the value of the tuning curve at each speed was computed by averaging the values larger than 4/5 of the peak value of each response time trace.

To quantify tuning curves using a single numerical metric and facilitate comparisons across conditions, we calculated the center of mass of the curve using the formula:

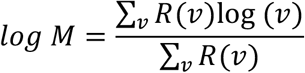

where *R*(*v*) was the computed response value at a specific velocity *v. R*(*v*) was set to be 0 if the response values were negative, making sure the geometric mean of the velocities were weighted by non-negative responses. This yields the log velocity of the tuning curve weighted by the response values.

### Numerical modeling

#### Linear-nonlinear (LN) model to test effects of sparsity changes on a simple model

We constructed a linear-nonlinear (LN) model (**Fig. S1A**) to test if our filter measurements were biased by changing the sparsity of the stimulus ^54^. The model output *r*_*LN*_(*t*) is defined as:

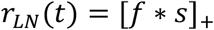

where the stimulus *s*(*t*) was linearly filtered by a high pass high-pass filter *f*(*t*) and then positively rectified. The high-pass filter was of the form:

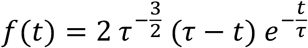

with *τ* = 80 *ms*.

#### Nonlinear-linear (NL) model for initial impulse response analysis

Extracting the light and dark impulse responses separately to represent the temporal processing properties of a neuron is related to modelling it using a nonlinear-linear (NL) model (**Fig. S1C**). To explore how such responses behave when analyzed by the techniques in our paper, we created a simple model with output *r*_*NL*_(*t*) defined as:

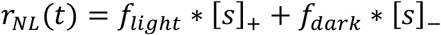

where [·]_+_ and [·]_−_ denote positive and negative rectification respectively, by which stimulus *s*(*t*) was separated into light and dark parts and linearly filtered by two distinct filters:

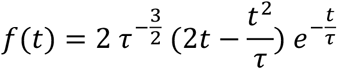

With *τ* = 100 ms for *f*_*light*_(*t*) and *τ* = 200 ms for *f*_*dark*_(*t*). These two filters are bandpass filters that integrate to 0. We computed *r*_*NL*_(*t*) in response to the same sparse and dense ternary noise stimuli used in our experiments and obtained filters and light and dark impulse responses to compare with the original filters *f*_*light*_ and *f*_*dark*_ (**Fig. S1D, E**). Importantly, when dense stimuli are used, every stimulus location that is not used to compute the light impulse response is used to compute the dark impulse response. This divides all snippets of the response into two categories, those associated with light and dark frames, while the sum of the snippets at each delay after the light or dark frame must equal the mean response. This imposes that the light and dark impulse responses to dense stimuli have equal and opposite deviations from the mean response, regardless of how the response was created.

To explore this symmetry imposed by the dense stimulus, we computed the model response to a uniform noise stimulus, which replaces each light/dark frame in the preceding ternary noise stimulus with a contrast drawn from a uniform distribution between –1 and 1. Then we extracted the impulse responses corresponding to various contrast bins (**Fig. S1F, G**). These different impulse responses do not have the symmetry imposed by the binary, dense stimulus, as discussed above, and instead produce two impulse responses with different timescales to the light and dark flashes. In contrast, the neural responses to the light and dark frames have similar timecourses and opposite deviations from the mean, suggesting that the close-to-linear responses to dense stimuli do not stem solely from the symmetry imposed by the dense stimulus.

### Biophysical plausible linear-nonlinear-linear (LNL) model for sparsity adaptation

We constructed a linear-nonlinear-linear (LNL) model to account for the observed phenomenon of sparsity adaptation. The model output *r*_*LNL*_(*t*) is defined as:

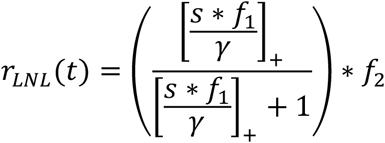

In this functional form, the stimulus *s*(*t*) was first filtered by a high-pass linear filter *f*_1_(*t*):

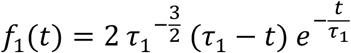

with *τ*_1_ = 80 *ms*. Then the output was positively rectified and scaled by dividing by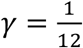 . Then a static nonlinearity function 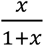 was applied before the output was filtered by a second low-pass filter *f*_2_(*t*):

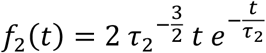

with *τ*_2_ = 150 ms. Both filters are normalized so that their *L*_2_-norm is equal to 1. And the scaling factor γ tuned the degree of saturation in the model.

### Quantification and statistical analysis

Each individual fly was treated as an independent measurement for statistical analysis. The ROI responses within a single fly were averaged together to generate each fly’s mean response. In the figures, solid lines along with shaded error bars represent the mean ± SEM of responses across all flies. N values in figure legends denote the number of individual flies analyzed. Significance throughout was assessed throughout using relatively conservative nonparametric Wilcoxon signed-rank tests (unpaired data) and Wilcoxon rank-sum tests (paired data).

## Declaration of generative AI and AI-assisted technologies in the writing process

During the preparation of this work, the first author used Chat-GPT only to improve grammar while drafting the initial prose. After using this tool, the authors have reviewed and edited the content through many subsequent drafts and take full responsibility for the content of the publication.

## Notes

### Competing Interest Statement

The authors have declared no competing interest.

## Citations

1. Weber, A.I., Krishnamurthy, K., and Fairhall, A.L. (2019). Coding Principles in Adaptation. Annual Review of Vision Science, Vol 5 5, 427–449.

2. Cafaro, J. (2016). Multiple sites of adaptation lead to contrast encoding in the Drosophila olfactory system. Physiol Rep 4.

3. Gorur-Shandilya, S., Demir, M., Long, J., Clark, D.A., and Emonet, T. (2017). Olfactory receptor neurons use gain control and complementary kinetics to encode intermittent odorant stimuli. Elife 6.

4. Cao, L.H., Jing, B.Y., Yang, D., Zeng, X., Shen, Y., Tu, Y., and Luo, D.G. (2016). Distinct signaling of Drosophila chemoreceptors in olfactory sensory neurons. Proc Natl Acad Sci U S A 113, E902–911.

5. Reisert, J., and Matthews, H.R. (1999). Adaptation of the odour-induced response in frog olfactory receptor cells. J Physiol 519 Pt 3, 801–813.

6. Wen, B., Wang, G.I., Dean, I., and Delgutte, B. (2009). Dynamic Range Adaptation to Sound Level Statistics in the Auditory Nerve. Journal of Neuroscience 29, 13797–13808.

7. Laughlin, S.B., and Hardie, R.C. (1978). Common Strategies for Light Adaptation in the Peripheral Visual Systems of Fly and Dragonfly. Journal of Comparative Physiology 128, 319–340.

8. Burkhardt, D.A. (1994). Light Adaptation and Photopigment Bleaching in Cone Photoreceptors in-Situ in the Retina of the Turtle. Journal of Neuroscience 14, 1091–1105.

9. Nikonov, S.S., Kholodenko, R., Lem, J., and Pugh, E.N. (2006). Physiological features of the S- and M-cone photoreceptors of wild-type mice from single-cell recordings. Journal of General Physiology 127, 359–374.

10. Juusola, M., Kouvalainen, E., Jarvilehto, M., and Weckstrom, M. (1994). Contrast Gain, Signal-to-Noise Ratio, and Linearity in Light-Adapted Blowfly Photoreceptors. Journal of General Physiology 104, 593–621.

11. Nagel, K.I., and Doupe, A.J. (2006). Temporal processing and adaptation in the songbird auditory forebrain. Neuron 51, 845–859.

12. Cooke, J.E., King, A.J., Willmore, B.D.B., and Schnupp, J.W.H. (2018). Contrast gain control in mouse auditory cortex. J Neurophysiol 120, 1872–1884.

13. Maravall, M., Petersen, R.S., Fairhall, A.L., Arabzadeh, E., and Diamond, M.E. (2007). Shifts in coding properties and maintenance of information transmission during adaptation in barrel cortex. PLoS Biol 5, e19.

14. Baccus, S.A., and Meister, M. (2002). Fast and slow contrast adaptation in retinal circuitry. Neuron 36, 909–919.

15. Rieke, F. (2001). Temporal contrast adaptation in salamander bipolar cells. J Neurosci 21, 9445–9454.

16. Brenner, N., Bialek, W., and de Ruyter van Steveninck, R. (2000). Adaptive rescaling maximizes information transmission. Neuron 26, 695–702.

17. Srinivasan, M.V., Laughlin, S.B., and Dubs, A. (1982). Predictive coding: a fresh view of inhibition in the retina. Proc R Soc Lond B Biol Sci 216, 427–459.

18. Laughlin, S. (1981). A simple coding procedure enhances a neuron’s information capacity. Z Naturforsch C Biosci 36, 910–912.

19. Vanhateren, J.H. (1992). Theoretical Predictions of Spatiotemporal Receptive-Fields of Fly Lmcs, and Experimental Validation. Journal of Comparative Physiology a-Neuroethology Sensory Neural and Behavioral Physiology 171, 157–170.

20. Drews, M.S., Leonhardt, A., Pirogova, N., Richter, F.G., Schuetzenberger, A., Braun, L., Serbe, E., and Borst, A. (2020). Dynamic Signal Compression for Robust Motion Vision in Flies. Curr Biol 30, 209–221 e208.

21. Matulis, C.A., Chen, J., Gonzalez-Suarez, A.D., Behnia, R., and Clark, D.A. (2020). Heterogeneous Temporal Contrast Adaptation in Drosophila Direction-Selective Circuits. Curr Biol 30, 222–236 e226.

22. Bonin, V., Mante, V., and Carandini, M. (2006). The statistical computation underlying contrast gain control. J Neurosci 26, 6346–6353.

23. Tkacik, G., Ghosh, A., Schneidman, E., and Segev, R. (2014). Adaptation to changes in higher-order stimulus statistics in the salamander retina. PLoS One 9, e85841.

24. Mechler, F., Reich, D.S., and Victor, J.D. (2002). Detection and discrimination of relative spatial phase by V1 neurons. J Neurosci 22, 6129–6157.

25. Felsen, G., Touryan, J., Han, F., and Dan, Y. (2005). Cortical sensitivity to visual features in natural scenes. PLoS Biol 3, e342.

26. Howard, J., Dubs, A., and Payne, R. (1984). The Dynamics of Phototransduction in Insects - a Comparative-Study. Journal of Comparative Physiology 154, 707–718.

27. Leutscher-Hazelhoff, J.T. (1975). Linear and non-linear performance of transducer and pupil in Calliphora retinula cells. J Physiol 246, 333–350.

28. Matthews, H.R., Fain, G.L., Murphy, R.L., and Lamb, T.D. (1990). Light adaptation in cone photoreceptors of the salamander: a role for cytoplasmic calcium. J Physiol 420, 447–469.

29. Normann, R.A., and Anderton, P.J. (1983). The incremental sensitivity curve of turtle cone photoreceptors. Vision Res 23, 1731–1733.

30. Enroth-Cugell, C., and Lennie, P. (1975). The control of retinal ganglion cell discharge by receptive field surrounds. J Physiol 247, 551–578.

31. Enroth-Cugell, C., and Robson, J.G. (1966). The contrast sensitivity of retinal ganglion cells of the cat. J Physiol 187, 517–552.

32. Grimes, W.N., Schwartz, G.W., and Rieke, F. (2014). The synaptic and circuit mechanisms underlying a change in spatial encoding in the retina. Neuron 82, 460–473.

33. Tikidji-Hamburyan, A., Reinhard, K., Seitter, H., Hovhannisyan, A., Procyk, C.A., Allen, A.E., Schenk, M., Lucas, R.J., and Munch, T.A. (2015). Retinal output changes qualitatively with every change in ambient illuminance. Nat Neurosci 18, 66–74.

34. Shapley, R.M., and Victor, J.D. (1978). The effect of contrast on the transfer properties of cat retinal ganglion cells. J Physiol 285, 275–298.

35. Smirnakis, S.M., Berry, M.J., Warland, D.K., Bialek, W., and Meister, M. (1997). Adaptation of retinal processing to image contrast and spatial scale. Nature 386, 69–73.

36. Chander, D., and Chichilnisky, E.J. (2001). Adaptation to temporal contrast in primate and salamander retina. J Neurosci 21, 9904–9916.

37. Hosoya, T., Baccus, S.A., and Meister, M. (2005). Dynamic predictive coding by the retina. Nature 436, 71–77.

38. Donoho, D.L. (2006). Compressed sensing. Ieee Transactions on Information Theory 52, 1289–1306.

39. Wallace, G.K. (1991). The Jpeg Still Picture Compression Standard. Communications of the Acm 34, 30–44.

40. Olshausen, B.A., and Field, D.J. (2004). Sparse coding of sensory inputs. Curr Opin Neurobiol 14, 481–487.

41. Froudarakis, E., Berens, P., Ecker, A.S., Cotton, R.J., Sinz, F.H., Yatsenko, D., Saggau, P., Bethge, M., and Tolias, A.S. (2014). Population code in mouse V1 facilitates readout of natural scenes through increased sparseness. Nature Neuroscience 17, 851–857.

42. Olshausen, B.A., and Field, D.J. (1996). Emergence of simple-cell receptive field properties by learning a sparse code for natural images. Nature 381, 607–609.

43. Yang, H.H., and Clandinin, T.R. (2018). Elementary Motion Detection in Drosophila: Algorithms and Mechanisms. Annu Rev Vis Sci 4, 143–163.

44. Meyer, H.G., Schwegmann, A., Lindemann, J.P., and Egelhaaf, M. (2014). Panoramic high dynamic range images in diverse environments.

45. Zonoobi, D., Kassim, A.A., and Venkatesh, Y.V. (2011). Gini Index as Sparsity Measure for Signal Reconstruction from Compressive Samples. Ieee Journal of Selected Topics in Signal Processing 5, 927–932.

46. Strother, J.A., Nern, A., and Reiser, M.B. (2014). Direct Observation of ON and OFF Pathways in the Drosophila Visual System. Current Biology 24, 976–983.

47. Behnia, R., Clark, D.A., Carter, A.G., Clandinin, T.R., and Desplan, C. (2014). Processing properties of ON and OFF pathways for Drosophila motion detection. Nature 512, 427–430.

48. Arenz, A., Drews, M.S., Richter, F.G., Ammer, G., and Borst, A. (2017). The Temporal Tuning of the Drosophila Motion Detectors Is Determined by the Dynamics of Their Input Elements. Curr Biol 27, 929–944.

49. Shinomiya, K., Huang, G., Lu, Z.Y., Parag, T., Xu, C.S., Aniceto, R., Ansari, N., Cheatham, N., Lauchie, S., Neace, E., et al. (2019). Comparisons between the ON- and OFF-edge motion pathways in the brain. Elife 8.

50. Maisak, M.S., Haag, J., Ammer, G., Serbe, E., Meier, M., Leonhardt, A., Schilling, T., Bahl, A., Rubin, G.M., Nern, A., et al. (2013). A directional tuning map of Drosophila elementary motion detectors. Nature 500, 212–216.

51. Creamer, M.S., Mano, O., and Clark, D.A. (2018). Visual Control of Walking Speed in Drosophila. Neuron 100, 1460–1473 e1466.

52. Chen, T.W., Wardill, T.J., Sun, Y., Pulver, S.R., Renninger, S.L., Baohan, A., Schreiter, E.R., Kerr, R.A., Orger, M.B., Jayaraman, V., et al. (2013). Ultrasensitive fluorescent proteins for imaging neuronal activity. Nature 499, 295-+.

53. Strother, J.A., Wu, S.T., Wong, A.M., Nern, A., Rogers, E.M., Le, J.Q., Rubin, G.M., and Reiser, M.B. (2017). The Emergence of Directional Selectivity in the Visual Motion Pathway of Drosophila. Neuron 94, 168–182 e110.

54. Chichilnisky, E.J. (2001). A simple white noise analysis of neuronal light responses. Network-Computation in Neural Systems 12, 199–213.

55. Yang, H.H., St-Pierre, F., Sun, X.L., Ding, X.Z., Lin, M.Z., and Clandinin, T.R. (2016). Subcellular Imaging of Voltage and Calcium Signals Reveals Neural Processing In Vivo. Cell 166, 245–257.

56. Kim, K.J., and Rieke, F. (2001). Temporal contrast adaptation in the input and output signals of salamander retinal ganglion cells. J Neurosci 21, 287–299.

57. Dana, H., Sun, Y., Mohar, B., Hulse, B.K., Kerlin, A.M., Hasseman, J.P., Tsegaye, G., Tsang, A., Wong, A., Patel, R., et al. (2019). High-performance calcium sensors for imaging activity in neuronal populations and microcompartments. Nature Methods 16, 649-+.

58. Nemenman, I. (2012). Gain control in molecular information processing: lessons from neuroscience. Physical Biology 9.

59. Clark, D.A., Benichou, R., Meister, M., and da Silveira, R.A. (2013). Dynamical Adaptation in Photoreceptors. Plos Computational Biology 9.

60. Borst, A., Flanagin, V.L., and Sompolinsky, H. (2005). Adaptation without parameter change: Dynamic gain control in motion detection. Proceedings of the National Academy of Sciences of the United States of America 102, 6172–6176.

61. Klapoetke, N.C., Murata, Y., Kim, S.S., Pulver, S.R., Birdsey-Benson, A., Cho, Y.K., Morimoto, T.K., Chuong, A.S., Carpenter, E.J., Tian, Z., et al. (2014). Independent optical excitation of distinct neural populations. Nat Methods 11, 338–346.

62. Jin, L., Han, Z., Platisa, J., Wooltorton, J.R.A., Cohen, L.B., and Pieribone, V.A. (2012). Single Action Potentials and Subthreshold Electrical Events Imaged in Neurons with a Fluorescent Protein Voltage Probe. Neuron 75, 779–785.

63. Mano, O., Creamer, M.S., Matulis, C.A., Salazar-Gatzimas, E., Chen, J.Y., Zavatone-Veth, J.A., and Clark, D.A. (2019). Using slow frame rate imaging to extract fast receptive fields. Nature Communications 10.

64. Takemura, S.Y., Bharioke, A., Lu, Z., Nern, A., Vitaladevuni, S., Rivlin, P.K., Katz, W.T., Olbris, D.J., Plaza, S.M., Winston, P., et al. (2013). A visual motion detection circuit suggested by Drosophila connectomics. Nature 500, 175–181.

65. Takemura, S.Y., Nern, A., Chklovskii, D.B., Scheffer, L.K., Rubin, G.M., and Meinertzhagen, I.A. (2017). The comprehensive connectome of a neural substrate for ‘ON’ motion detection in Drosophila. Elife 6.

66. Ketkar, M.D., Sporar, K., Gur, B., Ramos-Traslosheros, G., Seifert, M., and Silies, M. (2020). Luminance Information Is Required for the Accurate Estimation of Contrast in Rapidly Changing Visual Contexts. Curr Biol 30, 657–669 e654.

67. Clark, D.A., Bursztyn, L., Horowitz, M.A., Schnitzer, M.J., and Clandinin, T.R. (2011). Defining the Computational Structure of the Motion Detector in. Neuron 70, 1165–1177.

68. Davis, F.P., Nern, A., Picard, S., Reiser, M.B., Rubin, G.M., Eddy, S.R., and Henry, G.L. (2020). A genetic, genomic, and computational resource for exploring neural circuit function. Elife 9.

69. Nern, A., Pfeiffer, B.D., and Rubin, G.M. (2015). Optimized tools for multicolor stochastic labeling reveal diverse stereotyped cell arrangements in the fly visual system. Proceedings of the National Academy of Sciences of the United States of America 112, E2967–E2976.

70. Jing, M., Li, Y.X., Zeng, J.Z., Huang, P.C., Skirzewski, M., Kljakic, O., Peng, W.L., Qian, T.R., Tan, K., Zou, J., et al. (2020). An optimized acetylcholine sensor for monitoring in vivo cholinergic activity. Nature Methods 17, 1139-+.

71. Serbe, E., Meier, M., Leonhardt, A., and Borst, A. (2016). Comprehensive Characterization of the Major Presynaptic Elements to the Drosophila OFF Motion Detector. Neuron 89, 829–841.

72. Freifeld, L., Clark, D.A., Schnitzer, M.J., Horowitz, M.A., and Clandinin, T.R. (2013). GABAergic lateral interactions tune the early stages of visual processing in Drosophila. Neuron 78, 1075–1089.

73. Adelson, E.H., and Bergen, J.R. (1985). Spatiotemporal energy models for the perception of motion. J Opt Soc Am A 2, 284–299.

74. Hassenstein, B., and Reichardt, W. (1956). Systemtheoretische analyse der zeit-, reihenfolgen-und vorzeichenauswertung bei der bewegungsperzeption des rüsselkäfers chlorophanus. Zeitschrift für Naturforschung B 11, 513–524.

75. Gonzalez-Suarez, A.D., Zavatone-Veth, J.A., Chen, J., Matulis, C.A., Badwan, B.A., and Clark, D.A. (2022). Excitatory and inhibitory neural dynamics jointly tune motion detection. Curr Biol 32, 3659–3675 e3658.

76. Gruntman, E., Reimers, P., Romani, S., and Reiser, M.B. (2021). Non-preferred contrast responses in the Drosophila motion pathways reveal a receptive field structure that explains a common visual illusion. Current Biology 31, 5286-+.

77. Kim, A.J., Fenk, L.M., Lyu, C., and Maimon, G. (2017). Quantitative Predictions Orchestrate Visual Signaling in Drosophila. Cell 168, 280–294 e212.

78. Haikala, V., Joesch, M., Borst, A., and Mauss, A.S. (2013). Optogenetic control of fly optomotor responses. J Neurosci 33, 13927–13934.

79. Hu, Q., and Victor, J.D. (2010). A set of high-order spatiotemporal stimuli that elicit motion and reverse-phi percepts. Journal of Vision 10.

80. Chen, J.Y., Mandel, H.B., Fitzgerald, J.E., and Clark, D.A. (2019). Asymmetric ON-OFF processing of visual motion cancels variability induced by the structure of natural scenes. Elife 8.

81. Clark, D.A., Fitzgerald, J.E., Ales, J.M., Gohl, D.M., Silies, M.A., Norcia, A.M., and Clandinin, T.R. (2014). Flies and humans share a motion estimation strategy that exploits natural scene statistics. Nature Neuroscience 17, 296–303.

82. Fitzgerald, J.E., Katsov, A.Y., Clandinin, T.R., and Schnitzer, M.J. (2011). Symmetries in stimulus statistics shape the form of visual motion estimators. Proc Natl Acad Sci U S A 108, 12909–12914.

83. Dror, R.O., O’Carroll, D.C., and Laughlin, S.B. (2001). Accuracy of velocity estimation by Reichardt correlators. J Opt Soc Am A Opt Image Sci Vis 18, 241–252.

84. Lin, A.C., Bygrave, A.M., de Calignon, A., Lee, T., and Miesenbock, G. (2014). Sparse, decorrelated odor coding in the mushroom body enhances learned odor discrimination. Nat Neurosci 17, 559–568.

85. Schiller, P.H. (1992). The ON and OFF channels of the visual system. Trends Neurosci 15, 86–92.

86. Suzuki, H., Thiele, T.R., Faumont, S., Ezcurra, M., Lockery, S.R., and Schafer, W.R. (2008). Functional asymmetry in Caenorhabditis elegans taste neurons and its computational role in chemotaxis. Nature 454, 114–117.

87. Scholl, B., Gao, X., and Wehr, M. (2010). Nonoverlapping sets of synapses drive on responses and off responses in auditory cortex. Neuron 65, 412–421.

88. Laughlin, S.B., van Steveninck, R.R.D., and Anderson, J.C. (1998). The metabolic cost of neural information. Nature Neuroscience 1, 36–41.

89. Sterling, P., and Freed, M. (2007). How robust is a neural circuit? Visual neuroscience 24, 563–571.

90. Gjorgjieva, J., Sompolinsky, H., and Meister, M. (2014). Benefits of pathway splitting in sensory coding. J Neurosci 34, 12127–12144.

91. Kohn, J.R., Portes, J.P., Christenson, M.P., Abbott, L.F., and Behnia, R. (2021). Flexible filtering by neural inputs supports motion computation across states and stimuli. Curr Biol 31, 5249–5260 e5245.

92. Geffen, M.N., de Vries, S.E.J., and Meister, M. (2007). Retinal ganglion cells can rapidly change polarity from off to on. Plos Biology 5, 640–650.

93. Desimone, R., and Duncan, J. (1995). Neural mechanisms of selective visual attention. Annu Rev Neurosci 18, 193–222.

94. Maunsell, J.H.R. (2015). Neuronal Mechanisms of Visual Attention. Annu Rev Vis Sci 1, 373–391.

95. Maimon, G. (2011). Modulation of visual physiology by behavioral state in monkeys, mice, and flies. Curr Opin Neurobiol 21, 559–564.

96. Pinto, Y., van der Leij, A.R., Sligte, I.G., Lamme, V.A., and Scholte, H.S. (2013). Bottom-up and top-down attention are independent. J Vis 13, 16.

97. Connor, C.E., Egeth, H.E., and Yantis, S. (2004). Visual attention: bottom-up versus top-down. Curr Biol 14, R850–852.

98. Koch, C., and Ullman, S. (1985). Shifts in selective visual attention: towards the underlying neural circuitry. Hum Neurobiol 4, 219–227.

99. Itti, L., Koch, C., and Niebur, E. (1998). A model of saliency-based visual attention for rapid scene analysis. Ieee Transactions on Pattern Analysis and Machine Intelligence 20, 1254–1259.

100. Itti, L., and Koch, C. (2000). A saliency-based search mechanism for overt and covert shifts of visual attention. Vision Research 40, 1489–1506.

101. Knierim, J.J., and van Essen, D.C. (1992). Neuronal responses to static texture patterns in area V1 of the alert macaque monkey. J Neurophysiol 67, 961–980.

102. Egeth, H.E., and Yantis, S. (1997). Visual attention: control, representation, and time course. Annu Rev Psychol 48, 269–297.

103. Itti, L., and Koch, C. (2001). Computational modelling of visual attention. Nat Rev Neurosci 2, 194–203.

104. Posner, M.I. (1980). Orienting of attention. Q J Exp Psychol 32, 3–25.

105. Schreij, D., Owens, C., and Theeuwes, J. (2008). Abrupt onsets capture attention independent of top-down control settings. Percept Psychophys 70, 208–218.

106. Wang, F., Chen, M., Yan, Y., Zhaoping, L., and Li, W. (2015). Modulation of Neuronal Responses by Exogenous Attention in Macaque Primary Visual Cortex. J Neurosci 35, 13419–13429.

107. Klapoetke, N.C., Nern, A., Peek, M.Y., Rogers, E.M., Breads, P., Rubin, G.M., Reiser, M.B., and Card, G.M. (2017). Ultra-selective looming detection from radial motion opponency. Nature 551, 237-+.

108. Tanaka, R., and Clark, D.A. (2022). Neural mechanisms to exploit positional geometry for collision avoidance. Current Biology 32, 2357-+.

109. Keles, M.F., and Frye, M.A. (2017). Object-Detecting Neurons in Drosophila. Curr Biol 27, 680–687.

110. Tanaka, R., and Clark, D.A. (2020). Object-Displacement-Sensitive Visual Neurons Drive Freezing in Drosophila. Curr Biol 30, 2532–2550 e2538.

111. Marvin, J.S., Borghuis, B.G., Tian, L., Cichon, J., Harnett, M.T., Akerboom, J., Gordus, A., Renninger, S.L., Chen, T.W., Bargmann, C.I., et al. (2013). An optimized fluorescent probe for visualizing glutamate neurotransmission. Nature Methods 10, 162–170.

112. Rister, J., Pauls, D., Schnell, B., Ting, C.Y., Lee, C.H., Sinakevitch, I., Morante, J., Strausfeld, N.J., Ito, K., and Heisenberg, M. (2007). Dissection of the peripheral motion channel in the visual system of Drosophila melanogaster. Neuron 56, 155–170.

113. Silies, M., Gohl, D.M., Fisher, Y.E., Freifeld, L., Clark, D.A., and Clandinin, T.R. (2013). Modular Use of Peripheral Input Channels Tunes Motion-Detecting Circuitry. Neuron 79, 111–127.

114. Tuthill, J.C., Nern, A., Holtz, S.L., Rubin, G.M., and Reiser, M.B. (2013). Contributions of the 12 Neuron Classes in the Fly Lamina to Motion Vision. Neuron 79, 128–140.

115. Jenett, A., Rubin, G.M., Ngo, T.T.B., Shepherd, D., Murphy, C., Dionne, H., Pfeiffer, B.D., Cavallaro, A., Hall, D., Jeter, J., et al. (2012). A GAL4-Driver Line Resource for Neurobiology. Cell Reports 2, 991–1001.

116. Isaacman-Beck, J., Paik, K.C., Wienecke, C.F.R., Yang, H.H., Fisher, Y.E., Wang, I.E., Ishida, I.G., Maimon, G., Wilson, R.I., and Clandinin, T.R. (2020). SPARC enables genetic manipulation of precise proportions of cells. Nature Neuroscience 23, 1168-+.

117. Schilling, T., and Borst, A. (2015). Local motion detectors are required for the computation of expansion flow-fields. Biology Open 4, 1105–1108.

118. Brainard, D.H. (1997). The psychophysics toolbox. Spatial Vision 10, 433–436.

119. Kleiner, M., Brainard, D., and Pelli, D. (2007). What’s new in Psychtoolbox-3? Perception 36, 14–14.

120. Pelli, D.G. (1997). The VideoToolbox software for visual psychophysics: Transforming numbers into movies. Spatial Vision 10, 437–442.

121. Pologruto, T.A., Sabatini, B.L., and Svoboda, K. (2003). ScanImage: Flexible software for operating laser scanning microscopes. Biomedical Engineering Online 2.

122. Hurley, N., and Rickard, S. (2009). Comparing Measures of Sparsity. Ieee Transactions on Information Theory 55, 4723–4741.

123. Antoni, J. (2016). The infogram: Entropic evidence of the signature of repetitive transients. Mechanical Systems and Signal Processing 74, 73–94.

124. de Vries, S.E.J., and Clandinin, T. (2013). Optogenetic Stimulation of Escape Behavior in Drosophila melanogaster. Jove-Journal of Visualized Experiments.

125. Creamer, M.S., Mano, O., Tanaka, R., and Clark, D.A. (2019). A flexible geometry for panoramic visual and optogenetic stimulation during behavior and physiology. Journal of Neuroscience Methods 323, 48–55.

126. Wilson, R.I., Turner, G.C., and Laurent, G. (2004). Transformation of olfactory representations in the Drosophila antennal lobe. Science 303, 366–370.

127. Meyer, F. (1994). Topographic Distance and Watershed Lines. Signal Processing 38, 113–125.

128. Salazar-Gatzimas, E., Chen, J., Creamer, M.S., Mano, O., Mandel, H.B., Matulis, C.A., Pottackal, J., and Clark, D.A. (2016). Direct Measurement of Correlation Responses in Drosophila Elementary Motion Detectors Reveals Fast Timescale Tuning. Neuron 92, 227–239.

129. Salazar-Gatzimas, E., Agrochao, M., Fitzgerald, J.E., and Clark, D.A. (2018). The Neuronal Basis of an Illusory Motion Percept Is Explained by Decorrelation of Parallel Motion Pathways. Current Biology 28, 3748-+.

